# Alternative splicing contributes to plasticity and regulatory divergence in locally adapted house mice from the Americas

**DOI:** 10.1101/2025.07.03.662996

**Authors:** Megan Phifer-Rixey, Joseph R. Ward, Katya L. Mack

## Abstract

Alternative splicing is a major driver of transcriptome and proteome variation, but the role of alternative splicing in regulatory evolution remains understudied. Alternative splicing can also contribute to phenotypic plasticity, which may be critical when taxa colonize new environments. Here, we investigate variation in alternative splicing among new wild-derived strains of mice from different climates in the Americas on both a standard and high-fat diet. We show that alternative splicing is widespread and highly context-dependent. Comparisons between strains on different diets revealed abundant gene-by-environment interactions affecting alternative splicing, with most genes showing strain- and sex-specific diet responses. More often than not, genes that were differentially spliced between strains were not differentially expressed, adding to evidence that the two regulatory mechanisms often act independently. Moreover, differentially spliced genes were more widely expressed across tissues but also less central to biological networks than differentially expressed genes, suggesting differences in pleiotropic constraint. Importantly, divergence in alternative splicing was found to be predominantly driven by *cis-* regulatory changes. However, *trans* changes affecting splicing make be central to plasticity as they were impacted more by environmental variation. Finally, we performed scans for selection and found that, while genes with splicing divergence more often co-localized with genomic outliers associated with metabolic traits, they were not enriched for genomic outliers. Overall, our results provide evidence that alternative splicing plays an important role in gene regulation in house mice, contributing to adaptation and plasticity.

## Introduction

The regulation of gene expression plays a key role in adaptive evolution and divergence (Mack and Nachman 2017; Signor and Nuzhdin 2018). While the importance of gene regulation in adaptive evolution is increasingly appreciated (Stern and Orgogozo 2008; Fraser 2013; O’Brown et al. 2015; Verta and Jones 2019; Mack et al. 2023), the majority of studies have focused exclusively on changes in transcript abundance, often loosely referred to as “gene expression”. However, other mechanisms of gene regulation are common in eukaryotic lineages and can contribute to phenotypic differences within and between species (Schaefke et al. 2018; Verta and Jacobs 2022; C.J. Wright et al. 2022). Alternative splicing regulates gene expression by enabling the production of multiple unique mRNA transcripts from a single gene (i.e., isoforms). In humans and mice, the vast majority of genes have multiple isoforms (Pan et al. 2008; Tian et al. 2021). Moreover, alternative splicing can result in nonsense transcripts and thus can not only contribute to protein diversity but also to the regulation of transcript abundance (C.J. Wright et al. 2022). Alternative splicing can evolve independently of gene expression and is often controlled by different loci (Li et al. 2016; Qi et al. 2022). Consequently, alternative splicing and gene expression can act in complementary or conflicting ways to contribute to phenotypic evolution (Singh et al. 2017; Jacobs and Elmer 2021). Alternative splicing has now been implicated in ecologically significant traits in several systems (e.g., (Tovar-Corona et al. 2015; Howes et al. 2017; Mallarino et al. 2017; Smith et al. 2018)) and some comparative studies suggest that alternative splicing may even evolve more quickly than changes in gene expression, thereby potentiating rapid evolution (Barbosa-Morais et al. 2012; Merkin et al. 2012).

Alternative splicing may also be an important molecular mechanism contributing to phenotypic plasticity (Verta and Jacobs 2022; C.J. Wright et al. 2022). Phenotypic plasticity, where a genotype produces different phenotypes in response to environmental variation, plays a central role in mediating how organisms respond to changes in their local environment (Scheiner 1993; West-Eberhard, M.J. 2003; Sommer 2020). Phenotypic plasticity can evolve when genotypes vary in their response to environmental perturbations (genotype-by-environment interactions, or “GxE”), contributing to adaptive divergence when plastic responses increase fitness (Price et al. 2003; Lande 2009; Walter et al. 2022). Studies of plasticity and GxE interactions often focus on differences in transcript abundance associated with environment variation (e.g., (Campbell-Staton et al. 2021; Ballinger et al. 2023; Harry-Paul et al. 2024; Siddiq et al. 2024; Mack et al. 2025)). Alternative splicing, as a source of transcriptional variation, presents another avenue for phenotypic plasticity to evolve (Marden 2008). By increasing the diversity of the proteome, alternative splicing may allow organisms to respond rapidly to environmental changes or stressors. This may be particularly critical in seasonal environments, where large but predictable fluctuations in resources or climatic factors may lead to strong selection on plastic responses (Steward et al. 2022). For example, alternative splicing has been shown to play a significant role in temperature buffering in some systems (Healy and Schulte 2019; Martin Anduaga et al. 2019). Nevertheless, the role of alternative splicing in driving plastic responses to novel environments and in phenotypic adaptation remains largely underexplored (Singh and Ahi 2022).

The rapid expansion of house mice (*Mus musculus domesticus*) in the Americas presents a valuable opportunity to investigate the role of alternative splicing in plasticity and adaptation. House mice are human commensals and have recently colonized the Americas. Within the last ∼500 years, house mice have moved into diverse habitats and climates, accompanied by changes in body size, morphology, physiology, and behavior (Lynch 1992; Phifer-Rixey et al. 2018; Dumont et al. 2024). One particularly interesting example is changes in body mass, which vary with latitude. Mice from higher latitudes in the Americas are significantly larger, potentially as a thermoregulatory adaptation to different climates (i.e., Bergman’s rule)(Lynch 1992; Phifer-Rixey et al. 2018). Scans for selection in these populations have identified primarily non-coding variants, suggesting that genetic variation that affects gene regulation is a major driver of body size evolution (Phifer-Rixey et al. 2018; Gutiérrez-Guerrero et al. 2024). Supporting this idea, many of the genes highlighted in scans for selection also show differences in gene expression (Mack et al. 2018; Phifer-Rixey et al. 2018; Ballinger et al. 2023; Mack et al. 2025). While previous work in this system has primarily focused on transcript abundance (Mack et al. 2018; Ballinger et al. 2023; Durkin et al. 2024), the regulation of alternative splicing may also play an important role in evolved and plastic responses between populations of house mice. A recent study found evidence of splicing variation that differed in frequency among populations of wild mice from North America, indicating either evolved and/or plastic differences in splicing contribute to variation in these populations (Manahan and Nachman 2024). Differences in alternative splicing have also been implicated in diet-induced obesity and the regulation of energy metabolism in mice and humans (Vernia et al. 2016; Chao et al. 2021; Nomura et al. 2024), motivating the study of differences in alternative splicing in the context of adaptive body size evolution.

Importantly, there is growing evidence that plastic responses to environmental variation may play a significant role in phenotypic variation and adaptive divergence in house mice from the Americas (Bittner et al. 2021; Ballinger et al. 2023; Mack et al. 2025). Body mass is a complex and highly plastic trait, known to be affected by environmental factors, including diet. In humans and mice, genotype-by-diet interactions affecting body weight are common (Loos and Bouchard 2008; Bachmann et al. 2022; K.M. Wright et al. 2022). Variation in dietary content and seasonal differences in access to food can serve as a strong selection pressure, for both constitutive differences and plastic responses between populations. We recently demonstrated that diet can play an important role in body mass variation and transcriptional divergence in mice from the Americas (Mack et al. 2025). Using new wild derived strains originating from different climates and crosses among them, we characterized body size and patterns of transcript abundance on either a standard or a high-fat diet. Genotype, diet, and GxE interactions were all found to influence body weight, with widespread condition-specific gene regulation underlying gene expression differences between mice from different locations (Mack et al. 2025). These findings underscore the importance of environmental variation and GxE interactions in shaping complex trait variation, particularly for traits involved in environmental adaptation.

Here, we build on those results to investigate the role of alternative splicing in plasticity and adaptive divergence. Leveraging the same experimental framework, we ask how genetic and environmental variation, in this case diet, contribute to differences in alternative splicing among wild-derived inbred strains originating from 4 localities across the Americas. First, we characterized the extent and magnitude of alternative splicing differences between strains, sexes, and diet treatments, finding that alternative splicing was common, but highly context-dependent. Next, as alternative splicing and transcript abundance can evolve independently, we asked whether the same or different genes show differences in relation to strain and diet. We found that there was not a strong signal of shared regulation. In most cases, genes that were differentially spliced between strains were not also differentially expressed. Moreover, while both differentially expressed and differentially spliced genes were more pleiotropic than other genes, differentially spliced genes tended to be more widely expressed across tissues than differentially expressed genes but were less central to gene networks based on protein-protein interactions. These differences in aspects of pleiotropy may reflect differences in evolutionary constraint on these regulatory mechanisms. Using crosses between mice across a latitudinal gradient, we then investigated how *cis* and *trans* changes contributed to differences in alternative splicing, and how regulatory divergence is affected by diet. We found that splicing divergence was largely driven by *cis-*regulatory changes, identifying a modest number of *cis*-by-diet interactions impacting splicing divergence. However, *trans* changes affecting splicing were more impacted by diet and thus may be central to plasticity. Finally, we found that differentially spliced genes were not enriched for genomic outliers from selection scans, but they did tend to co-localize with genomic outliers associated with metabolic traits. Together, our results underscore the importance of alternative splicing as a mechanism contributing to regulatory adaptation and plasticity.

## Results and Discussion

### Variation in alternative splicing and transcript levels in mice from the Americas

House mice from different geographic regions in the Americas have been shown to differ in aspects of body size, metabolism, and morphology (Phifer-Rixey et al. 2018; Dumont et al. 2024), with mice from higher latitudes showing higher average body mass (Lynch 1992; Phifer-Rixey et al. 2018). Here, we leverage the results of a recent study focusing on transcript abundance and body size to investigate the role of alternative splicing in plasticity and adaptation in this system (Mack et al. 2025). Briefly, male and female mice from wild-derived inbred strains originating from localities across the Americas were fed either a standard or high-fat diet for twelve weeks post-weaning (Figure 1A, B; Mack et al. 2025). Body size measures were collected regularly and, at the end of the experiment, liver tissue was collected for transcriptome sequencing from five strains derived from four populations (Figure 1A; Mack et al. 2025). We reanalyzed the RNA-seq data, using the program rMATS, to search for five major types of splice events (1) skipped exon, (2) intron retention, (3) mutually exclusive exons, (4) alternative 3′ splice site, and (5) alternative 5′ splice site (Figure 1C)(Shen et al. 2014; Wang et al. 2024). We detected 174,645 alternative splicing events across genotypes and diets. This corresponded to 16,775 multi-exonic genes expressed in our dataset. The most common event type detected in our data was exon skipping (∼73% of events), followed by mutually exclusive exons (∼17% of events). In a principal component analysis of alternative splicing events, samples clustered by inbred strain of origin on the first (7.9% of variation) and second principal components (PC)(5.97% of variance)(Fig 1D). Strain origin was also a major axis of variation for transcript levels (PC2, 21% of variance)(Mack et al. 2025). Strains from more northern latitudes (Edmonton, Saratoga Springs) and more southern latitudes (Gainesville, Manaus) were separated on PC2. However, in contrast to analysis of transcript levels in which samples were clearly separated by sex on PC2 (Mack et al. 2025), in our analysis of splicing variation, samples did not cluster strongly by sex.

**Figure 1.**
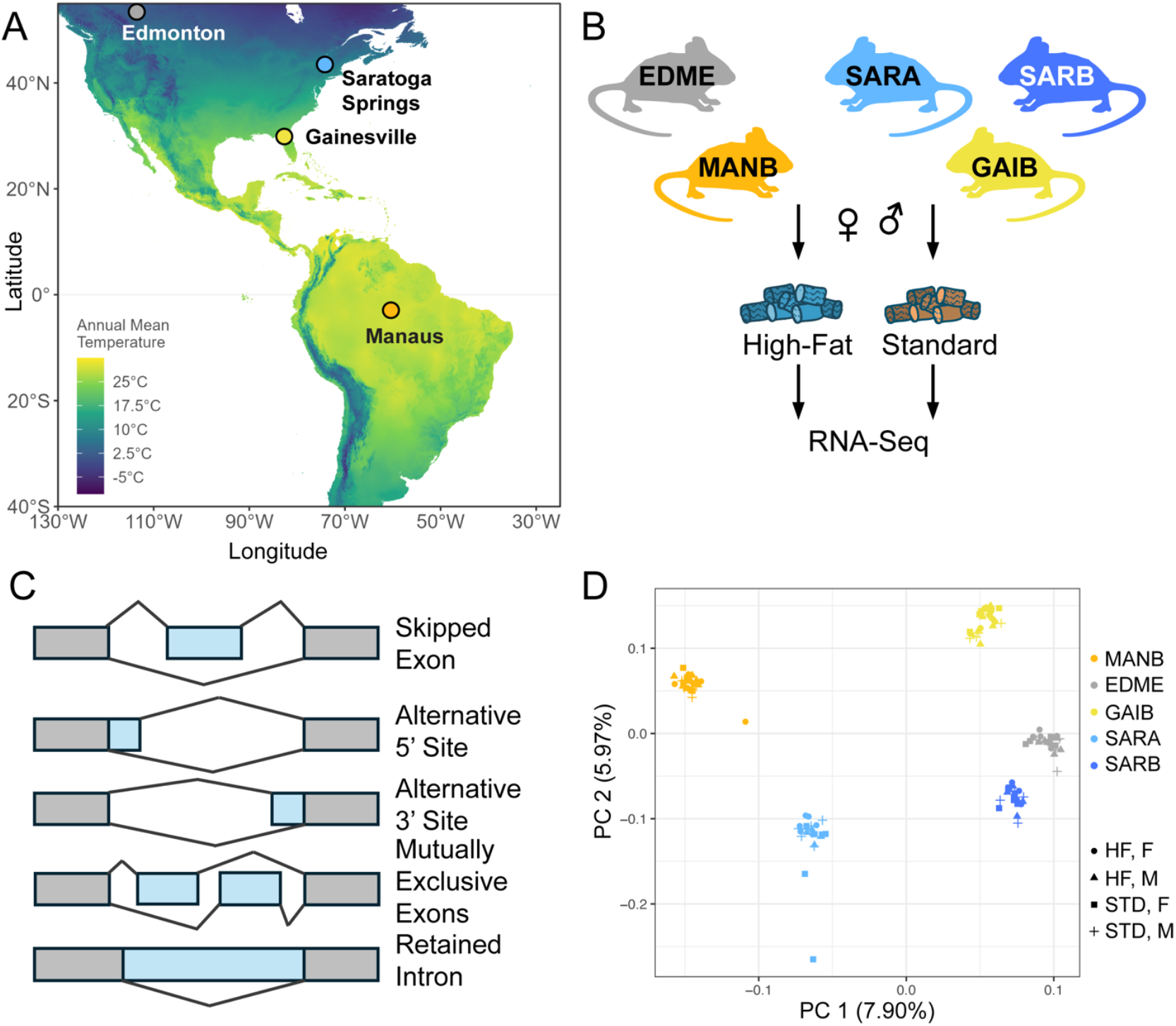
Alternative splicing variation in wild-derived house mouse strains from the Americas. **A**. Locations from which each strain was derived: EDME: Edmonton, Alberta, Canada; GAIB: Gainesville, Florida, USA; MANB: Manaus, Amazonas, Brazil; SARA, SARB: Saratoga Springs, New York, USA. **B**. Males and females from each strain were reared on either a high-fat (HF) or standard diet (STD). Liver tissue was collected from each mouse for RNA-seq. **C**. Five alternative splicing event types were surveyed across strains. **D**. Principal component analysis of splicing variation showed that strains cluster by locality on PC 1 and PC2.

### Extensive alternative splicing divergence between strains from divergent climates

Pairwise comparisons identified 2,084-3,672 differential splicing events between strains (FDR<0.05) on a standard diet among 5,556 genes. Approximately 42% of genes were differentially spliced between at least two strains. Among differential splicing events, exon skipping remained the most prevalent (8,247 events total, 3,920 genes), followed by mutually exclusive exons (3,550 events, 1,857 genes). The greatest number of differential splicing events were observed between females from the Gainesville (GAIB) and Edmonton (EDME) strains (3,672 events) and from the Gainesville (GAIB) and Manaus (MANB) strains (3,621 events), where the fewest were observed between males from the Saratoga Springs strains (SARA and SARB), likely reflecting their shared population origin (2,084 events).

As low abundance splicing events may represent splicing mistakes, we also examined differential splicing at a threshold of difference in percent spliced in (ΔPSI) of >10% (Grantham and Brisson 2018)(Table S1). After this filtering, 3,197 genes (24%) were differentially spliced between at least two strains, indicating many splicing differences between strains were of small magnitude despite being statistically significant. As in the other analysis, the majority of these events were exon skipping (4,165 events) or mutually exclusive exons (1,548).

### Alternative splicing across diet types is highly strain- and sex-specific

Characterizing the contribution of alternative splicing to environmental plasticity is important for understanding the adaptive potential of alternative splicing in the colonization of new environments. To this end, we asked how diet affected alternative splicing and whether diet-dependent effects were strain-specific. Data from mice on either a standard or high-fat diet were compared to identify changes in splicing associated with diet. We discovered 3,376 differential splicing events across 2,153 genes associated with diet, corresponding to approximately 13% of genes surveyed. This was similar to the number of genes identified as differentially expressed in response to diet (2,163 genes, 10% of genes surveyed; Mack et al. 2025). Approximately 1,071 of these genes were also significant at a >10% ΔPSI threshold (see above), indicating many splicing events identified were of a smaller magnitude. Most diet-related differences in splicing were strain-specific (Figure 2A). No genes showed diet-related differential splicing across all strains/sexes, and only 34% of differentially spliced genes were observed across multiple genotypes. Intron retention differences between diets were more often shared across multiple genotypes relative to other splicing events (19% of shared events) (Fisher’s exact test, *P*=0.0006). We also observed some strain-specific patterns in the frequency of splicing events (Figure 2B, Table S2). For example, females from the Edmonton strain (EDME) showed a higher proportion of intron retention differences between diets than other lines (Fisher’s exact test, 9.12 x10^−48^). Females from the New York SARA line showed a higher rate of mutually exclusive exons (Fisher’s exact test, 2.31 × 10^−27^).

**Figure 2.**
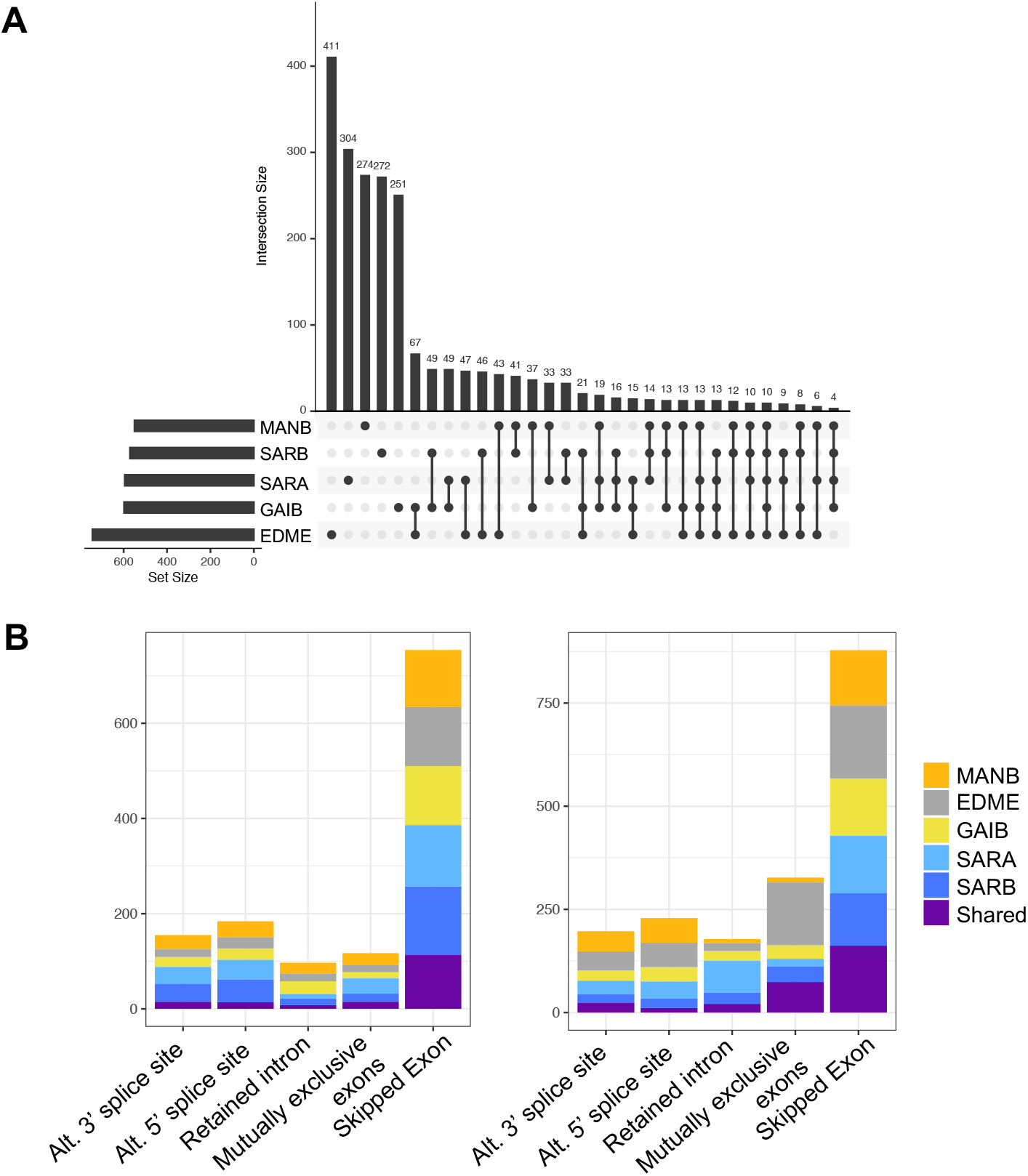
Alternative splicing differences associated with diet. **A**. UpSet plot showing the overlap in differentially spliced genes across strains between a high-fat and standard diet. Alternative splicing was strain-specific for the majority of genes. **B**. Event types for differential spliced genes across diets for each strain. The “shared” bar represents genes that showed differential splicing in more than one strain.

Notably, in contrast to overall variation in splicing, diet-associated differential splicing events were nearly always sex-specific (92% of significant events). More alternative splicing differences related to diet were observed in females compared to males. In females, we identified 2,111 differential splicing events (1,483 genes) and in males we identified 1,526 (1,118 genes). This was consistent with patterns of differential expression, with females having a greater proportion of genes with differential expression across diets (1,862 vs. 571 genes in males and females, respectively (Mack et al. 2025)).

### Sex-biased splicing is highly context-specific

Sex-specific splicing can play an important role in sexual dimorphism and sex-specific adaptation (Blekhman et al. 2010; Rogers et al. 2020; García-Pérez et al. 2023). To investigate sex-specific differences in splicing in our dataset, we compared splicing events between males and females. We identified sex-biased splicing at 1,738 and 1,598 genes between males and females on a standard and high-fat diet, respectively. This corresponded to ∼3%-6% of genes surveyed per strain. Surprisingly, genes with evidence for differential splicing between the sexes were largely unique to one diet type (Table S3), with only 7-8% of sex-biased splicing events being shared across diet contexts. For example, a total of 846 genes showed sex-biased splicing in the SARA strain, but only 89 genes were sex-biased under both diets. This indicates that sex-specific differences in splicing can be highly context-dependent. Comparing across event types, we found that mutually exclusive splicing of exons was shared across diet contexts more often than other splicing events (15% vs. 6% shared; Fisher’s exact test, *P*=1.42 × 10^−16^).

Sex-biased splicing was also largely strain-specific (Figure S1). Most genes showed sex-biased splicing in only one strain (∼77% in standard fat and 81% in high-fat). Mutually exclusive splicing of exon events were again shared more often across strains than other splicing events (Fisher’s exact tests, Standard: *P*=3.34 x10^−5^ [21% vs.11% shared], High-fat: *P*=0.0015 [16% vs. 10% shared]). Only 15 and 12 genes were identified as differentially spliced between sexes across all strains under a standard and high-fat diet, respectively (Table S4). Only 5 genes show sex-specific splicing across all strains and both diet treatments (*Zbtb20, Scp2, Sult2a8, AI182371, Rpl22l1*). Overall, these results indicate that sexual dimorphism in alternative splicing is highly dependent on genetic background and environmental context.

### Non-redundant roles for gene expression and alternative splicing in divergence and diet responses

An important open question is whether alternative splicing plays a complementary or contrasting role in expression evolution, targeting either the same or independent genes or pathways (Jacobs and Elmer 2021; Verta and Jacobs 2022; Innes et al. 2024). To address this question, we compared the number and identity of genes that showed differences in mRNA abundance (e.g., gene expression) and alternative splicing. The majority of differentially spliced genes did not show differences in gene expression. Examining pairwise differences between strains on a standard diet, we found an overlap of ∼7-10% of genes between differential splicing and differential expression. The overlap between genes was more than expected by chance (Fisher’s exact tests, *P-*values<1.66 x10^−14^). However, differences in expression and splicing between strains are more likely to be detected for highly expressed genes due to increased power, which may bias this approach. Indeed, mean expression for overlapping genes was higher than for genes with only differential splicing or expression between strains (Permutation tests, 10,000 permutations: *P*=0.048 and *P*<0.0001, respectively). To explore this relationship further, we binned genes by mean expression and examined overlap for genes at different average expression levels for SARB and MANB females, the comparison that showed the highest overlap in the original analysis. More overlap than expected by chance was observed for all bins (Fisher’s exact tests, *P*<0.024), with the exception of the bin containing genes with lowest average expression (*P*=0.56)(see Methods), suggesting that expression level alone was not driving this pattern (Table S5).

We then compared genes with evidence for differential splicing and differential expression in response to diet. Of differentially spliced genes in response to diet, a relatively small number also showed expression differences associated with diet in the same strain (1-39 genes per genotype, 0.4-10% of genes). Looking across strains, 307 genes showed both differences in splicing and expression associated with diet, though these changes were most often not identified within the same strain and/or sex (228/307 genes). Genes identified as diet-responsive in both expression and splicing were enriched for GO ontology terms related lipid and fatty acid metabolism (e.g., lipid metabolic process, BH-corrected *p*-value (*q*)=3.19×10^−6^, long-chain fatty acid metabolic process, *q*=1.41 × 10-5) and Reactome pathways related to metabolism (e.g., Metabolism of lipids, *q*=1.62 x10^−4^) and absorption, distribution, metabolism, and excretion (ADME)(e.g., Atorvastatin ADME, *q*=3.53×10^−2^, Drug ADME, *q*=1.77×10^−2^). There were very few genes that showed both differential expression and splicing for more than one strain and sex (14 genes; i.e., *Gstm6, Gstm3, Gstm1, Fam107b, Cyp3a41b, Rad51b, Cyp3a41a, Pnldc1, Cyp3a44, Zfand4, Camk1d, Impg2, Ehhadh, Aprt*). However, this overlap included multiple members of the GSTM family, which are organized in a gene cluster on chromosome 3. Variation at these genes has been linked to susceptibility to environmental toxins and metabolic disease (Hu et al. 2024; Alnasser 2025). Low overlap between differentially expressed genes and differentially spliced genes within strains suggests that these two regulatory mechanisms are operating on different genes and/or pathways in response to diet.

### Increased pleiotropy of genes involved in diet responses and divergence

Pleiotropy, where a single locus influences multiple phenotypes, can constrain plastic and evolved responses. Alternative splicing provides a potential mechanism for reducing constraint on highly pleiotropic genes though the generation of alternative isoforms (Verta and Jacobs 2022). Relative to changes in gene expression, it has been hypothesized that splicing may be less evolutionarily constrained because it can shift proportions of isoforms without disrupting the expression of essential isoforms (C.J. Wright et al. 2022). We sought to investigate the relationship between plastic responses to diet and pleiotropy in house mice through three approaches. First, we examined developmental and tissue-specificity of genes using an expression atlas of 8 tissues across multiple developmental stages (Papatheodorou et al. 2018), as genes that are expressed in multiple tissues are more likely to affect multiple traits (Watanabe et al. 2019). Level of pleiotropy is associated with the number of protein-protein interactions, as well as the number of biological processes a gene is involved in (He and Zhang 2006). We investigated the number of protein-protein interactions associated with each gene using high confidence annotations from the STRING database for *Mus musculus*. To estimate the number of biological processes a gene was involved in, we annotated genes to GO ontology Biological Process terms (McGirr and Martin 2018; Jacobs et al. 2024).

Differentially expressed genes were found to expressed in more tissues/developmental stages (Permutation test [PT], 10,000 permutations, *P-*value<0.0001), were associated with more GO ontology terms (PT, *P*=0.001), and with a greater number of protein-protein interactions (PT, *P*<0.0001) than non-differentially expressed genes. Altogether, this suggests that genes that are differentially expressed in response to diet are potentially more pleiotropic and under greater evolutionary constraint. In contrast, we found that while differentially spliced genes were also expressed in a larger number of tissue-developmental stages than non-differentially spliced genes (PT, *P*<0.0001), these genes were not associated with a more ontology terms (PT, *P*=0.95), and with fewer protein-protein interactions (PT, *P*<0.0001). Comparing differentially expressed and differentially spliced genes directly, we also found that differentially expressed genes were associated with more protein-protein interactions than differentially spliced genes (PT, *P*<0.0001), but that differentially expressed genes were expressed in fewer tissues/developmental stages (PT, *P*<0.0001) (Figure S2). This suggests that differentially expressed genes have a higher level of tissue-specificity but may also be more central to biological networks than differentially spliced genes.

Next, to examine features of evolved differences, we compared differentially expressed and spliced genes between strains collected from higher and lower latitudes with divergent thermal environments (Saratoga Springs and Edmonton vs. Gainesville and Manaus; Figure 1). Results were similar to that of our analysis of plastic genes. Differentially expressed genes were associated with more protein-protein interactions (PT, *P*=0.027), more GO Biological Process terms (PT, *P*<0.0001), and expressed across more tissues (PT, *P*<0.0001) relative to genes that were not differentially expressed. Genes with differential splicing between strains were associated with fewer protein-protein interactions (PT, *P*<0.0001), but were expressed across more tissues on average (PT, *P*<0.0001) relative to non-differentially spliced genes. Comparing these directly, differentially spliced genes were expressed across more tissues on average (PT, *P*<0.0001) and had fewer protein-protein interactions (PT, *P*=0.01). Altogether, these results suggest that differentially spliced and differentially expressed genes are both potentially more pleiotropic than other genes but may differ in aspects of their evolutionary constraint based on network interactions and tissue specificity.

### *Cis-* and *trans-* gene regulatory divergence drive differences in alternative splicing between strains from divergent climates

Divergence in alternative splicing between populations can arise via changes in *cis* and/or *trans* regulation. Changes in *trans* regulation reflect evolution of RNA-binding proteins while changes in *cis* reflect evolution of binding sites in the transcripts. Distinguishing between the two mechanisms addresses fundamental evolutionary processes, including the potential impact of pleiotropy on the evolution of gene regulation (Wittkopp and Kalay 2011; Signor and Nuzhdin 2018). To identify regulatory divergence between mice from divergent climates due to changes in *cis*, we examined differences in splicing between alleles in F1 offspring between strains. In an F1 individual, pre-mRNAs are exposed to the same *trans*-acting environment. Consequently, allele-specific splicing differences will reflect differences at *cis-*regulatory elements (Wittkopp et al. 2004; McManus et al. 2014). Changes in the *trans*-acting environment can then be inferred by comparing differences between the parental strains with allele-specific differences in the F1 offspring (Wittkopp et al. 2004; McManus et al. 2014).

Mice originating from the cold temperate locality of Saratoga Springs, New York (SARA, SARB) were crossed with mice from two localities representing lower latitudes, Gainesville, Florida, USA (GAIB) and Manaus, Brazil (MANB) (Figure 1A) to produce F1s. As with the parental strains, F1s (GAIBxSARA or SARBxMANB) were reared on standard or high-fat diet (for details, see Mack et al. 2025). An average of 13,389,010 allele-specific paired-end reads were aligned for each sex/diet treatment. We restricted our analysis to allele-specific splicing events supported by at least 20 sequencing reads in F1 hybrids (see Methods). This allowed us to test for allele-specific alternative splicing (ASAS) at 3,760 genes for SARBxMANB hybrids. We identified 637 genes with evidence for ASAS, indicative of *cis-*regulatory divergence. In GAIBxSARA crosses, we identified 574 genes (of 3,807) with evidence of ASAS (Table S6). As in parental comparisons, the majority of events with evidence for ASAS were skipped exons.

To identify the relative contribution of *cis-* vs. *trans-*regulatory changes to divergence in alternative splicing between lines, we compared allele-specific splicing in hybrids with differential splicing between parental lines on a standard diet (McManus et al. 2014; Gao et al. 2015). We identified evidence for both *cis* and *trans* divergence contributing to differences in splicing between strains. However, consistent with previous studies (McManus et al. 2014; Gao et al. 2015), we found that splicing divergence was more often attributable to divergence in *cis*. Approximately ∼4% of events that could be tested showed solely *cis*-regulatory divergence with 0.4%-0.7% showing solely *trans* divergence in each cross (Figure 3)(Table S7).

**Figure 3.**
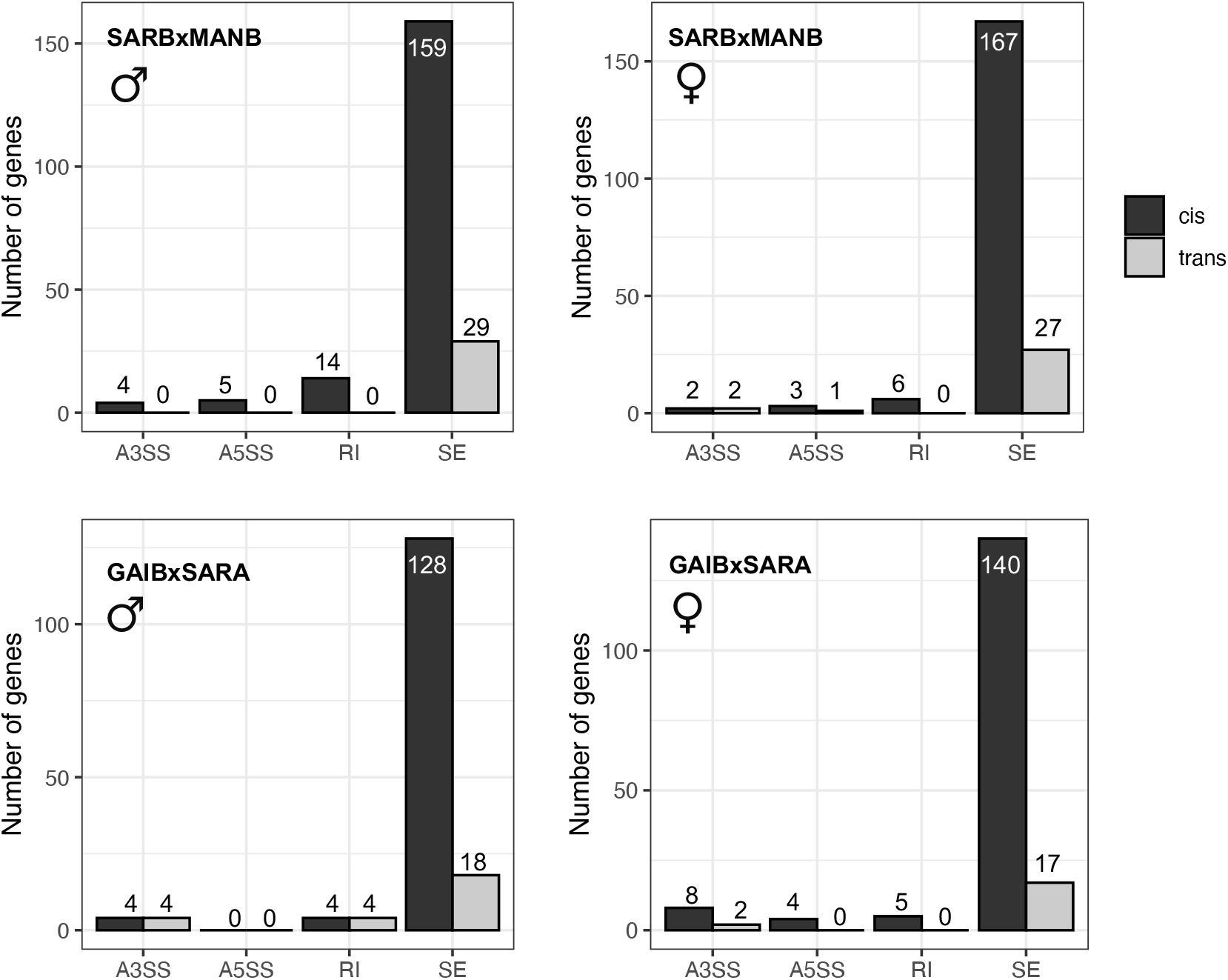
*Cis* and *trans* changes contribute to differences in splicing between strains from different climates. Bar charts below show the number of genes attributed to either *cis* (left bars) or *trans* (right bars) for each splicing event (A3SS: alternative 3’ splice site; A5SS: alternative 5’ splice site; RI: retained intron; SE: skipped exon).

### Diet-specific gene regulatory divergence between strains

Next, we examined the relationship between gene regulatory divergence and plasticity in alternative splicing. As the majority of differential alternative splicing was a consequence of *cis-* regulatory changes, we first examined the relationship between *cis-*regulatory variation and diet. Allele-specific ratios from F1s subjected to each diet treatment were compared to understand the role of *cis-*by-environment interactions in driving alternative splicing variation. Allele-specific measures were correlated between high-fat and standard diets, with stronger correlations seen in the SARBxMANB cross (Figure 4; SARBxMANB, Spearman’s *rho*=0.63-0.65; GAIBxSARA, Spearman’s *rho*=0.28-0.31, *P*<0.0001).

**Figure 4.**
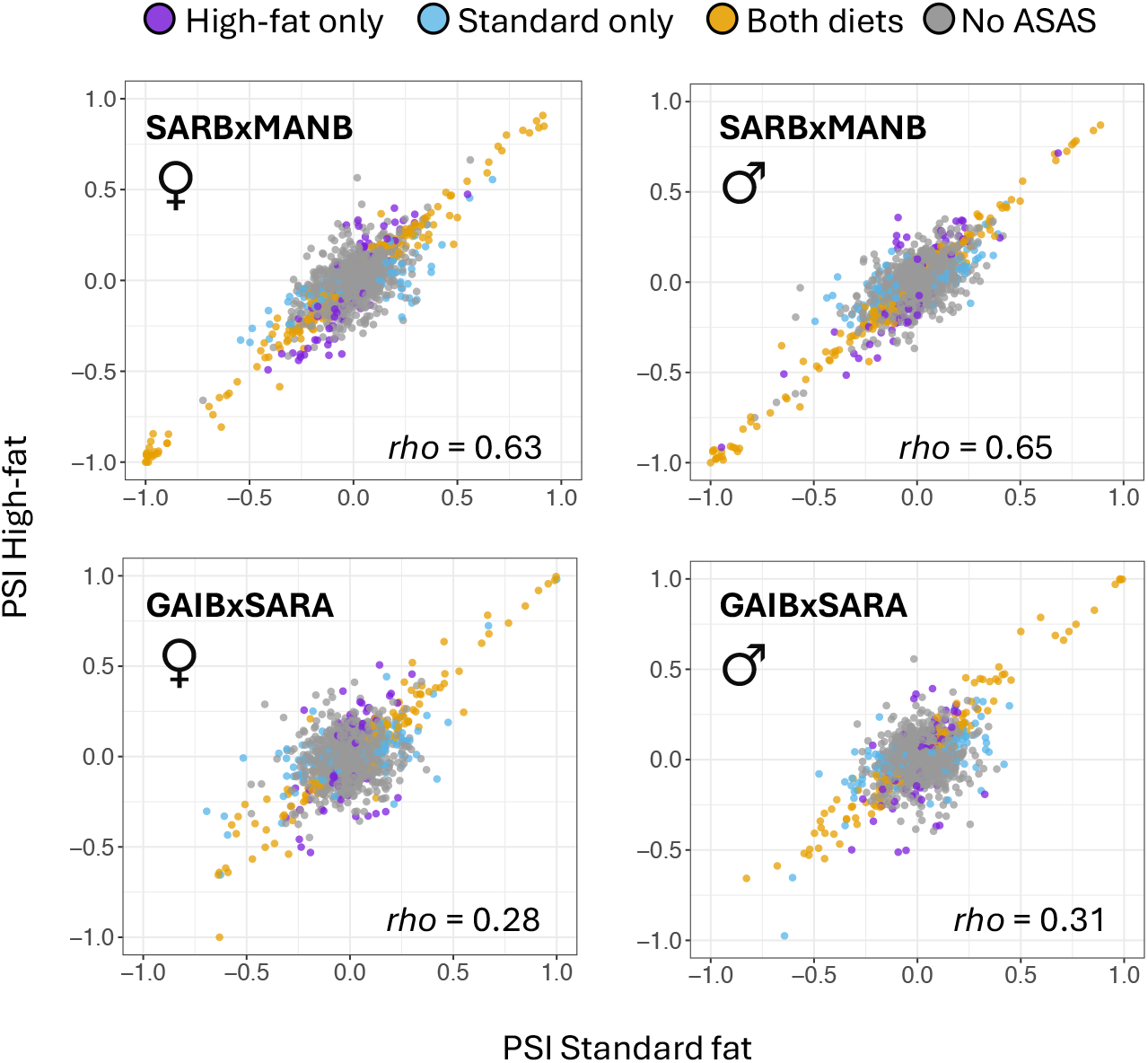
Evidence for diet-specific allele-specific alternative splicing (ASAS). Plots show comparisons of percent spliced in (PSI) proportions between alleles on the high-fat versus standard diet. Cases where ASAS is specific to one diet (purple and blue) highlight *cis*-by-diet interactions affecting alternative splicing.

Correlations between allele-specific measures across diets suggests *cis-*regulatory differences affecting splicing are often robust to environment. Consistent with this, many genes were identified as having allele-specific alternative splicing in both diets (39%-47% of genes with evidence for ASAS; Figure 4). This overlap was greater at higher |ΔPSI| thresholds (41-53% at 10% |ΔPSI| cut-off). Still, most genes were identified as having ASAS in only one diet in each comparison, revealing many diet-specific *cis*-regulatory effects (Figure 4). Examining the *trans* component, we also find *trans* effects on alternative splicing were correlated across diet treatments, though not as strongly as for *cis* effects (SARBxMANB, Spearman’s *rho*=0.49-0.50; GAIBxSARA, Spearman’s *rho*=0.8-0.11, *p*-values< 2.45 × 10^−11^). Using log_2_ fold changes as a proxy for effect size, we observed greater divergence in the *trans* component per gene between diets (Wilcoxon signed rank test, *P-*values<1.49 × 10^−33^). Together, these results suggest that *trans* differences affecting splicing are influenced more by environment than *cis* changes and thus may be more important in the evolution of plasticity.

Next, we formally tested for *cis*-by-diet interactions by comparing ΔPSI between alleles across the two diet treatments (see Methods). While power is limited to test for interactions, we identified 7 events with evidence for *cis-*by-diet effects on splicing (at FDR<0.1; Genes: *Trim24, C4bp, Hmgn1, Cyp4f15, Pip4p2, Rnf126, Tpk1*). *Trim24* is of particular interest due to its role in susceptibility to high-fat-induced hepatic steatosis (Wei et al. 2022). *TRIM24* plays an essential role in linking insulin signaling to processing bodies (Wei et al. 2022). Genes *C4bp* and *Hmgn1* have also been implicated in aspects of metabolism via mouse mutant models (Birling et al. 2021).

### Genomic outliers for climate and alternative splicing

Alternative splicing has been proposed as an important substrate for rapid adaptive evolution (Singh and Ahi 2022). If differences in alternative splicing contribute to adaptation, we might expect genes with evidence of alternative splicing to co-localize with signals of selection. Utilizing exome-sequence data collected from natural populations in North and South America (Phifer-Rixey et al. 2018; Ferris et al. 2021; Gutiérrez-Guerrero et al. 2024), we searched for genomic regions with signatures consistent with natural selection. Scans for differentiated regions were performed using a normalized version of the population branch statistic (*PBSn1*)(Yi et al. 2010; Malaspinas et al. 2016), which are commonly used to identify local selective sweeps (Shpak et al. 2025). We identified genomic outliers for each population by performing the test multiple times and varying the focal and outgroups, focusing on the most geographically distant populations (Northeastern USA vs. Manaus, Edmonton vs. Manaus, and Gainesville vs. Northeastern USA) (see Methods). We considered windows in the top 1% as genomic outliers in each comparison. We then compared genes with differential splicing between more northern latitudes (Northeastern USA, Edmonton) and more southern latitudes (Manaus, Gainesville) with outlier regions identified for each focal population (Table S8).

We identified 229 genes with differential splicing between southern and northern populations that overlapped a genomic outlier window (54-74 genes per population pair; Table S8). Genes with evidence for differential splicing that overlapped genomic outliers were enriched for mutant phenotypes related to metabolism relative to other genes expressed in the liver for which there was sufficient data to test for alternative splicing (Table S9). For example, genes with differential splicing between Edmonton and Manaus associated with outlier variants were enriched for the phenotype terms “abnormal postnatal growth/weight/body size” (MP:0002089), “abnormal body size” (MP:0003956), and “abnormal body composition” (MP:0005451)(FDR<0.05). However, genes with differential alternative splicing were not more likely to overlap outlier windows than other genes expressed in the liver (Chi square test, *P-values*>0.13). This suggests that, although splicing-related outliers are often related to metabolism, there is no evidence that loci associated with differences in splicing are enriched for signals of selection.

Genes harboring genomic outliers may be interesting candidates for phenotypic differences between strains, particularly when these genes are associated with metabolic phenotypes in mutant models. While we did not observe a significant association between differential alternative splicing and genomic outliers overall, we did identify a number of outlier genes with differential splicing and relevant mutant phenotypes and/or diet-interactions. One example of this was the gene *Akt2*, a genomic outlier in the Amazonas population. *Akt2* is an important regulator of lipid and glucose metabolism (Leavens et al. 2009). Another is *Mmp19*, a matrix metalloproteinase. Mice with deficient MMP19 exhibit higher body weights and gain more weight on a high-fat diet (Pendás et al. 2004; Molière et al. 2023).

We also compared our differential splicing results to that of a recent study of wild mice from a latitudinal transect in eastern North America (Florida to New Hampshire/Vermont)(Manahan and Nachman 2024). In that study, five genes were identified as candidates for adaptive splicing, based on the presence *cis*-regulatory variation associated with clinal patterns of splicing and overlapping signals of selection (*Mbl2, Pex26, Rnd2, Lasp1, Atg7;* (Manahan and Nachman 2024)). We also identified evidence for differential splicing between New York and Florida strains for 3 of these genes (*Mbl2, Pex26, and Atg7*). Of these, *Atg7* is of particular interest as a candidate gene for body size adaptation, as it plays a role in adipogenesis (Zhang et al. 2009) and obesity in mice (Ren et al. 2025). Isoforms of this gene have also previously been shown to play distinct roles in central energy metabolism (Ostacolo et al. 2024).

## Conclusions

Alternative splicing is a gene regulatory mechanism that has potential to contribute to plasticity and adaptation (Singh and Ahi 2022; Verta and Jacobs 2022; C.J. Wright et al. 2022). However, not only is the importance of alternative splicing in those processes not yet well established, but in most systems, there are still many fundamental open questions regarding the extent and drivers of variation in alternative splicing. Advances in analytical methods have made it possible to execute coordinated analysis of gene expression and alternative splicing from short-read data. While there are still some limitations of this approach, we now can characterize variation in alternative splicing and more directly compare both forms of gene regulation and their targets.

Here, we investigated the role of alternative splicing in plastic responses and transcriptional divergence in the context of the expansion of house mice into the Americas. Similar to what we previously found for differential expression across these strains (Mack et al. 2025), variation in alternative splicing was common and driven by both genetic and environmental variation.

Furthermore, alternative splicing differences associated with diet were most often strain-specific, providing strong evidence for gene-by-environment effects on splicing. The consistent signal of context dependence from both differential expression and differential splicing analyses highlights the importance of GxE interactions in shaping transcriptional variation and underscores the need to incorporate environmental and genetic variation into experimental investigations of gene regulation.

We also found commonalities in the genetic architecture of differential expression and differential splicing, but there were key differences as well. For example, both here for splicing and previously for gene expression (Mack et al. 2025), we found that *cis-*regulatory differences were more robust across treatments compared to changes in *trans*. Therefore, for both mechanisms, there is support for the hypothesis that *trans* differences are more responsive to environmental variation and thus may be critical in the evolution of plasticity. Also consistent with results for gene expression, we found that both *cis* and *trans* changes contributed to regulatory divergence in alternative splicing. However, for splicing, *cis*-regulatory changes greatly outnumbered changes in *trans* while contributions were more balanced between *cis-* and *trans*-regulatory changes involved in expression divergence (Mack et al. 2025). The central role of *cis*-regulation to splicing divergence is consistent with what has been observed in other studies, including between subspecies of house mice (Gao et al. 2015) and in fruit flies, humans, and *Arabidopsis* (McManus et al. 2014; Wang et al. 2019; García-Pérez et al. 2023).

The majority of genes for which we identified differential splicing were not differentially expressed, indicating that differential splicing introduces protein variation that is not captured by surveying gene expression differences alone. Differences in the genes impacted and the genetic architecture of alternative splicing and differential expression suggest that the two regulatory mechanisms evolve largely independently. While data are still relatively limited, this result echoes what has been seen in other systems, including salmonid fishes (Jacobs and Elmer 2021), butterflies (Steward et al. 2022), pea aphids (Grantham and Brisson 2018), and sunflowers (Innes et al. 2024), though greater overlap has also been observed (e.g., (Singh et al. 2017)).

Interestingly, we found that, unlike results for gene expression, sex was not a major driver of overall variation in alternative splicing, only in response to diet. This mirrors what has been observed for humans, where sex was not identified as a major driver of variation in splicing for most tissues (García-Pérez et al. 2023). This difference between expression and splicing suggests distinct roles for the two mechanisms in fundamental biological processes. Reinforcing this idea, we found that differentially expressed genes were associated with more protein-protein interactions but were expressed in fewer tissue/developmental stages compared to differentially spliced genes. Alternative splicing has been suggested as a mechanism for resolving evolutionary constraint at pleiotropic genes (Verta and Jacobs 2022), but our results point to broad differences in how pleiotropy may shape evolution and plasticity via these mechanisms that requires further investigation.

Importantly, while there is strong evidence that *cis*-regulatory variation impacting gene expression plays a central role in adaptive evolution in this system (Ballinger et al. 2023; Durkin et al. 2024; Mack et al. 2025), that was not the case for splicing. Genes with splicing changes were not enriched for signals of selection. Consequently, loci affecting transcript levels may be more relevant to rapid climate adaptation in house mice. Nevertheless, in both this study and a previous study of wild mice (Manahan and Nachman 2024), there were splicing-related outliers that are promising candidates for environmental adaptation. As splicing-related outliers were also found to be enriched for genes associated with metabolic phenotypes, splicing variation may be of particular interest in the context of body size evolution.

Our results add to a growing body of evidence that supports independent roles for regulation of transcript levels versus splicing in both plastic responses (Grantham and Brisson 2018; Steward et al. 2022) and evolved differences (Jacobs and Elmer 2021). While there are still many open questions, it is clear that incorporating alternative splicing can lead to new insights and contribute to a more comprehensive understanding of gene regulatory evolution.

## Methods

### Samples and Sequencing

Five newly derived strains of *M. m. domesticus* from the Americas were selected to evaluate the transcriptional response to high-fat diet: GAIB (Gainesville, FL), SARA and SARB (Sarasota Springs, NY), EDME (Edmonton, Alberta) and Brazil, MANB (Manaus, Amazonas). Strains were derived from brother-sister matings for at least 12 generations as previously described (Dumont et al. 2024). RNA extraction and library preparation proceeded as previously described (Mack et al. 2025). Briefly, RNA was extracted from frozen liver tissue and strand-specific libraries were prepared using NEBNext Ultra II RNA Library prep kits (New England Biolabs, Ipswich, MA, USA). Libraries were sequenced on the Illumina HiSeq instrument (2×150bp Paired End [PE] reads). To increase coverage, libraries were also sequenced using the same approach on the NovaSeq platform (2×150bp PE).

### Gene expression analysis

All sequencing data used are available online through NCBI BioProject (PRJNA1164275) (Mack et al. 2025). Gene expression analysis was performed as previously described (Mack et al. 2025). In short, reads were aligned with STAR (Dobin et al. 2013) to the *Mus musculus* reference genome (GRCm39) using the Ensembl GRCm39 annotation. To quantify gene expression, reads overlapping exons were counted using HTSeq (htseq-count function, default settings)(Anders et al. 2015). For principal component analysis (PCA), reads were transformed using variance stabilizing transformation in DESeq2 (Love et al. 2014). DESeq2 was then used to assess differential expression between strains, diet-treatments, and sex (Wald test). *P*-values were corrected with the Benjamini–Hochberg method. Genes with an adjusted *P-*value (FDR) of less than 0.05 were considered significantly differentially expressed between comparisons.

### Alternative splicing analysis

For alternative splicing analysis, reads were aligned with STAR in two-pass mode to the *Mus musculus* reference genome (GRCm39) using the Ensembl GRCm39 annotation. We used the event-based tool rMATs (version: turbo, 4.3.0)(Shen et al. 2014; Wang et al. 2024) to quantify splice junctions and test for differential splicing events across genotypes, sexes, and diet treatments. rMATS identified five splicing events: (1) alternative 3′ splice site, (2) skipped exon, (3) intron retention, (4) mutually exclusive exons and (5) alternative 5’ splice sites. Reads that align across splice junctions and within exons are used to estimate the “percent spliced in” (PSI) of each event for each sample.

Differential splicing is then calculated as the difference in PSI between comparisons (for example, for comparisons between the high-fat and standard diet for the SARB line, ΔPSI = mean(PSI_SARB high-fat_) - mean(PSI_SARB standard_)). ΔPSI ranges from 1 to −1, with the extreme values (1 and -1) representing fixed differences in the isoform proportions. A likelihood ratio test is used to test the significance of differential splicing events based on differential PSI values. We considered events significant at an adjusted *p*-values (FDR) of 0.05 based on supporting junction reads. We also evaluated splicing a more conservative threshold based on a ΔPSI cut-off of 10%. We report how this more conservative cut-off impacts our findings in the Results section and in Table S1, Table S6, and Table S8.

Genes with evidence for differential expression and differential splicing were compared at an FDR<0.05. To examine how expression level affects overlap, we removed lowly expressed genes (<20 average reads per sample) and then binned genes by average expression level using the R function “quartile”, with ∼20% of genes in each bin. Bins were then examined for overlap of differentially spliced and differentially expressed genes (Table S5).

### Examining features of differentially expressed and differentially spliced genes

We investigated pleiotropy via (1) protein-protein interactions, (2) Biological Process GO ontology terms, and (3) expression across tissues/developmental stages. Protein-Protein interactions were downloaded from the STRING database (Szklarczyk et al. 2023) and filtered for high confidence interactions (Scores >0.7; where scores range from 0-1, and 1 is the highest possible confidence). This threshold (>0.7) has previously been shown to capture relevant protein-protein interactions (von Mering et al. 2005). Protein annotations were annotated to Ensembl gene ids based on annotations downloaded from the STRING database (string-db.org). Genes were annotated to GO ontology terms downloaded from Mouse Genome Informatics and filtered for terms that were annotated as involved in Biological Process (Baldarelli et al. 2024).

To examine the presence or absence of expression across tissues and developmental stages, we downloaded expression data from Expression Atlas (Papatheodorou et al. 2018) for *Mus musculus domesticus* (European Nucleotide Archive Project PRJEB26869) comprised of 8 tissues at up to 14 developmental stages per tissue (94 tissue-timepoints total) ranging from embryonic day 10.5 to adulthood. A FPKM cut-off of 3 for a gene to be considered expressed for a given tissue-timepoint. We tested for differences in averages between sets using permutation tests, where labels for each group were shuffled 10,000 times. The means of the shuffled datasets were then compared to the observed mean to calculate a *p*-value.

### Allele-specific splicing

Variant calling for allele-specific assignment for the GAIBxSARA cross was performed as previously described (Mack et al. 2025). However, whole-genome sequencing data is available for SARA and MANB (Dumont et al. 2024). Therefore, variant calls for ASE for that cross were downloaded (https://doi.org/10.5281/zenodo.10728008) based on the whole-genome sequencing data and SNP calls produced (Dumont et al. 2024).

As with the parental strains, reads for F1 hybrids were mapped to the *Mus musculus* reference genome (GRCm39) in two-pass mode with STAR to identify allele-specific splicing (Dobin et al. 2013). We used the WASP implementation in STAR to mitigate mapping bias associated with the reference allele (Asiimwe and Alexander 2024). Reads overlapping an informative variant that passed WASP filtering were then retained to examine allele-specific splicing. Reads were then separated into bam files associated with their parental allele for analysis with rMATs. To increase confidence for estimates of allele-specific expression, we filtered junctions based on the number of supporting reads. Junctions supported by fewer than 20 reads in a replicate were discarded. We considered events significant at a Benjamini-Hochberg (BH) adjusted *p*-value (FDR) of 0.05.

### Gene Regulatory Assignment

To infer *cis-* vs. *trans*-regulatory divergence, the ratio of PSI values between parental alleles were compared to the allele-specific ratios in hybrids following the approach of Altman and Bland (Altman and Bland 2003) as previously described (McManus et al. 2014; Gao et al. 2015; Wang et al. 2019). Briefly, the PSI values between strains (e.g., SARB/MANB) were compared to allele-specific PSI ratios from hybrids (SARB allele/MANB allele). The standard error of differences in parental and allele-specific ratios was calculated and used to calculate Z-scores and *P*-values. *P-*values were then corrected using the BH procedure. Events with a significant difference between alleles in a hybrid, between parents, and no significant difference between ratios were considered divergent in *cis* alone. Events with a significant difference between parents, no significant difference in hybrids, and a significant difference in the ratios were considered evidence of *trans* only divergence (McManus et al. 2014). For comparisons in which sample size was unequal, individuals were randomly dropped to equalize sample size (5 individuals per comparison). We required at least 20 supporting reads per event per sample (Gao et al. 2015).The same approach was used to search for evidence of *cis*-by-diet interactions, i.e., comparing the allele-specific ratios from hybrids reared on a standard and high-fat diet (e.g., [high-fat SARB allele/high-fat MANB allele] vs. [standard SARB allele/standard MANB allele]).

### Scans for genomic outliers

To identify regions with signatures consistent with selection, we used previously generated exome data from North and South America (Phifer-Rixey et al. 2018; Gutiérrez-Guerrero et al. 2024) and whole-genome data from Eurasia (Harr et al. 2016). North and South American individuals were collected from the same geographic localities or localities near where inbred lines originate from (Phifer-Rixey et al. 2018; Dumont et al. 2024). Reads were mapped to *Mus musculus* reference genome (GRCm39) with BWA-MEM (Li and Durbin 2009) and variants were called with BCFtools (Danecek et al. 2021; Lefouili and Nam 2022) and then filtered for low quality calls (‘QUAL>20 && DP>100’).

We utilized a modification of the population branch statistic (PBS)(Yi et al. 2010) to perform a comparison between a focal population and the relevant thermally contrasting Americas population and a Eurasian outgroup (Germany) (*PBSn1*) (Malaspinas et al. 2016; Crawford et al. 2017). Population branch statistics estimate the branch lengths of focal populations relative to outgroups, as an approach for identifying loci with patterns consistent with local selective sweeps (Shpak et al. 2025). We restricted our SNP set to biallelic variants across the 3 populations being compared and required that at least six individuals in the focal branch be genotyped. VCFtools was used to calculate Weir and Cockerham *F*_*st*_ for each variant position between population pairs (Danecek et al. 2011). *PBSn1* scores were calculated for non-overlapping blocks of 5 SNPs as previously described (Ballinger et al. 2023).

The following comparisons were performed to identify genomic outliers for each population (focal population listed first and *in italics*):

1. *New Hampshire/Vermont, USA*; Manaus, Amazonas, Brazil; Germany
2. *Manaus, Amazonas, Brazil*; *New Hampshire/Vermont*, USA; Germany
3. *Gainesville, Florida, USA*; New Hampshire/Vermont, USA; Germany
4. *Edmonton, Canada*; Manaus, Amazonas, Brazil; Germany

We note that the New Hampshire/Vermont samples used in the PBS test are geographically close to the origin of the SARA and SARB line (Phifer-Rixey et al. 2018). We consider blocks in the top 1% of *PBSn1* scores outliers and do not attempt to assign *P* values to each SNP block (Malaspinas et al. 2016; Ballinger et al. 2023). These loci are considered “genomic outliers”, as population branch statistics do not consider neutral models of evolution. Genes overlapping SNP blocks were identified using the BEDtools “intersect” tool and Ensembl gene coordinates (GRCm39). The full list of splicing-related outliers can be found in Table S10.

### Enrichment analyses

Tests for enrichments of GO ontology terms and Reactome pathways were performed with PANTHER (Thomas et al. 2003). Tests for enrichments based on phenotypic associations in mice were performed with ModPhea (Weng and Liao 2017). Phenotype data is based on mutagenesis and knockdown experiments in mouse models available through MGI (Blake et al. 2009; Weng and Liao 2017).

## Data Availability

The sequencing data used in this project are available online through NCBI BioProject (PRJNA1164275).

## Acknowledgements

MPR is supported by NSF Division of Environmental Biology Award #2332998. KLM is funded by the National Institute of General Medical Sciences of the National Institutes of Health under Award Number P30 GM145499 and NIH R35GM154966. This work used Expanse at the San Deigo Supercomputer Center through allocation BIO230113 from the Advanced Cyberinfrastructure Coordination Ecosystem: Services & Support (ACCESS) program, which is supported by National Science Foundation grants #2138259, #2138286, #2138307, #2137603, and #2138296.

## References

Alnasser SM. 2025. The role of glutathione S-transferases in human disease pathogenesis and their current inhibitors. Genes & Diseases 12:101482.

Altman DG, Bland JM. 2003. Interaction revisited: the difference between two estimates. BMJ 326:219.

Anders S, Pyl PT, Huber W. 2015. HTSeq--a Python framework to work with high-throughput sequencing data. Bioinformatics 31:166–169.

Asiimwe R, Alexander D. 2024. STAR+WASP reduces reference bias in the allele-specific mapping of RNA-seq reads. bioRxiv:2024.01.21.576391.

Bachmann AM, Morel J-D, El Alam G, Rodríguez-López S, Imamura de lima T, Goeminne LJE, Benegiamo G, Loric S, Conti M, Sleiman MB, et al. 2022. Genetic background and sex control the outcome of high-fat diet feeding in mice. iScience 25:104468.

Baldarelli RM, Smith CL, Ringwald M, Richardson JE, Bult CJ, Mouse Genome Informatics Group. 2024. Mouse Genome Informatics: an integrated knowledgebase system for the laboratory mouse. Genetics 227:iyae031.

Ballinger MA, Mack KL, Durkin SM, Riddell EA, Nachman MW. 2023. Environmentally robust cis-regulatory changes underlie rapid climatic adaptation. Proceedings of the National Academy of Sciences 120:e2214614120.

Barbosa-Morais NL, Irimia M, Pan Q, Xiong HY, Gueroussov S, Lee LJ, Slobodeniuc V, Kutter C, Watt S, Çolak R, et al. 2012. The Evolutionary Landscape of Alternative Splicing in Vertebrate Species. Science 338:1587–1593.

Birling M-C, Yoshiki A, Adams DJ, Ayabe S, Beaudet AL, Bottomley J, Bradley A, Brown SD, Bürger A, Bushell W, et al. 2021. A resource of targeted mutant mouse lines for 5,061 genes. Nat Genet 53:416–419.

Bittner NKJ, Mack KL, Nachman MW. 2021. Gene expression plasticity and desert adaptation in house mice. Evolution 75:1477–1491.

Blake JA, Bult CJ, Eppig JT, Kadin JA, Richardson JE. 2009. The Mouse Genome Database genotypes::phenotypes. Nucleic Acids Res 37:D712–D719.

Blekhman R, Marioni JC, Zumbo P, Stephens M, Gilad Y. 2010. Sex-specific and lineage-specific alternative splicing in primates. Genome Res 20:180–189.

Campbell-Staton SC, Velotta JP, Winchell KM. 2021. Selection on adaptive and maladaptive gene expression plasticity during thermal adaptation to urban heat islands. Nat Commun 12:6195.

Chao Y, Jiang Y, Zhong M, Wei K, Hu C, Qin Y, Zuo Y, Yang L, Shen Z, Zou C. 2021. Regulatory roles and mechanisms of alternative RNA splicing in adipogenesis and human metabolic health. Cell Biosci 11:66.

Crawford JE, Alves JM, Palmer WJ, Day JP, Sylla M, Ramasamy R, Surendran SN, Black WC, Pain A, Jiggins FM. 2017. Population genomics reveals that an anthropophilic population of Aedes aegypti mosquitoes in West Africa recently gave rise to American and Asian populations of this major disease vector. BMC Biology 15:16.

Danecek P, Auton A, Abecasis G, Albers CA, Banks E, DePristo MA, Handsaker RE, Lunter G, Marth GT, Sherry ST, et al. 2011. The variant call format and VCFtools. Bioinformatics 27:2156–2158.

Danecek P, Bonfield JK, Liddle J, Marshall J, Ohan V, Pollard MO, Whitwham A, Keane T, McCarthy SA, Davies RM, et al. 2021. Twelve years of SAMtools and BCFtools. GigaScience 10:giab008.

Dobin A, Davis CA, Schlesinger F, Drenkow J, Zaleski C, Jha S, Batut P, Chaisson M, Gingeras TR. 2013. STAR: ultrafast universal RNA-seq aligner. Bioinformatics 29:15–21.

Dumont BL, Gatti DM, Ballinger MA, Lin D, Phifer-Rixey M, Sheehan MJ, Suzuki TA, Wooldridge LK, Frempong HO, Lawal RA, et al. 2024. Into the Wild: A novel wild-derived inbred strain resource expands the genomic and phenotypic diversity of laboratory mouse models. PLOS Genetics 20:e1011228.

Durkin SM, Ballinger MA, Nachman MW. 2024. Tissue-specific and cis-regulatory changes underlie parallel, adaptive gene expression evolution in house mice. PLOS Genetics 20:e1010892.

Ferris KG, Chavez AS, Suzuki TA, Beckman EJ, Phifer-Rixey M, Bi K, Nachman MW. 2021. The genomics of rapid climatic adaptation and parallel evolution in North American house mice. PLOS Genetics 17:e1009495.

Fraser HB. 2013. Gene expression drives local adaptation in humans. Genome Res. 23:1089–1096.

Gao Q, Sun W, Ballegeer M, Libert C, Chen W. 2015. Predominant contribution of cis-regulatory divergence in the evolution of mouse alternative splicing. Mol Syst Biol 11:816.

García-Pérez R, Ramirez JM, Ripoll-Cladellas A, Chazarra-Gil R, Oliveros W, Soldatkina O, Bosio M, Rognon PJ, Capella-Gutierrez S, Calvo M, et al. 2023. The landscape of expression and alternative splicing variation across human traits. Cell Genomics [Internet] 3. Available from: https://www.cell.com/cell-genomics/abstract/S2666-979X(22)00207-5

Grantham ME, Brisson JA. 2018. Extensive Differential Splicing Underlies Phenotypically Plastic Aphid Morphs. Molecular Biology and Evolution 35:1934–1946.

Gutiérrez-Guerrero YT, Phifer-Rixey M, Nachman MW. 2024. Across two continents: The genomic basis of environmental adaptation in house mice (Mus musculus domesticus) from the Americas. PLOS Genetics 20:e1011036.

Harr B, Karakoc E, Neme R, Teschke M, Pfeifle C, Pezer Ž, Babiker H, Linnenbrink M, Montero I, Scavetta R, et al. 2016. Genomic resources for wild populations of the house mouse, Mus musculus and its close relative Mus spretus. Sci Data 3:160075.

Harry-Paul YY, Lachapelle J, Ness RW. 2024. The Evolution of Gene Expression Plasticity During Adaptation to Salt in Chlamydomonas reinhardtii. Genome Biology and Evolution 16:evae214.

He X, Zhang J. 2006. Toward a Molecular Understanding of Pleiotropy. Genetics 173:1885–1891.

Healy TM, Schulte PM. 2019. Patterns of alternative splicing in response to cold acclimation in fish. Journal of Experimental Biology 222:jeb193516.

Howes TR, Summers BR, Kingsley DM. 2017. Dorsal spine evolution in threespine sticklebacks via a splicing change in MSX2A. BMC Biology 15:115.

Hu L, Yuan D, Zhu Q, Wu M, Tie M, Song S, Chen Y, Yang Y, He A. 2024. Evaluation of the role of hepatic Gstm4 in diet-induced obesity and dyslipidemia. Biochemical and Biophysical Research Communications 737:150920.

Innes PA, Goebl AM, Smith CCR, Rosenberger K, Kane NC. 2024. Gene expression and alternative splicing contribute to adaptive divergence of ecotypes. Heredity 132:120–132.

Jacobs A, Elmer KR. 2021. Alternative splicing and gene expression play contrasting roles in the parallel phenotypic evolution of a salmonid fish. Molecular Ecology 30:4955–4969.

Jacobs A, Velotta JP, Tigano A, Wilder AP, Baumann H, Therkildsen NO. 2024. Temperature-dependent gene regulatory divergence underlies local adaptation with gene flow in the Atlantic silverside. Evolution 78:1133–1149.

Lande R. 2009. Adaptation to an extraordinary environment by evolution of phenotypic plasticity and genetic assimilation. Journal of Evolutionary Biology 22:1435–1446.

Leavens KF, Easton RM, Shulman GI, Previs SF, Birnbaum MJ. 2009. Akt2 is required for hepatic lipid accumulation in models of insulin resistance. Cell Metab 10:405–418.

Lefouili M, Nam K. 2022. The evaluation of Bcftools mpileup and GATK HaplotypeCaller for variant calling in non-human species. Sci Rep 12:11331.

Li H, Durbin R. 2009. Fast and accurate short read alignment with Burrows–Wheeler transform. Bioinformatics 25:1754–1760.

Li YI, van de Geijn B, Raj A, Knowles DA, Petti AA, Golan D, Gilad Y, Pritchard JK. 2016. RNA splicing is a primary link between genetic variation and disease. Science 352:600–604.

Loos RJF, Bouchard C. 2008. FTO: the first gene contributing to common forms of human obesity. Obesity Reviews 9:246–250.

Love MI, Huber W, Anders S. 2014. Moderated estimation of fold change and dispersion for RNA-seq data with DESeq2. Genome Biology 15:550.

Lynch CB. 1992. Clinal Variation in Cold Adaptation in Mus domesticus: Verification of Predictions from Laboratory Populations. The American Naturalist 139:1219–1236.

Mack KL, Ballinger MA, Phifer-Rixey M, Nachman MW. 2018. Gene regulation underlies environmental adaptation in house mice. Genome Res. 28:1636–1645.

Mack KL, Landino NP, Tertyshnaia M, Longo TC, Vera SA, Crew LA, McDonald K, Phifer-Rixey M. 2025. Gene-by-environment interactions and adaptive body size variation in mice from the Americas. Molecular Biology and Evolution:msaf078.

Mack KL, Nachman MW. 2017. Gene Regulation and Speciation. Trends in Genetics 33:68–80.

Mack KL, Square TA, Zhao B, Miller CT, Fraser HB. 2023. Evolution of Spatial and Temporal cis-Regulatory Divergence in Sticklebacks. Molecular Biology and Evolution 40:msad034.

Malaspinas A-S, Westaway MC, Muller C, Sousa VC, Lao O, Alves I, Bergström A, Athanasiadis G, Cheng JY, Crawford JE, et al. 2016. A genomic history of Aboriginal Australia. Nature 538:207–214.

Mallarino R, Linden TA, Linnen CR, Hoekstra HE. 2017. The role of isoforms in the evolution of cryptic coloration in Peromyscus mice. Molecular Ecology 26:245–258.

Manahan DN, Nachman MW. 2024. Alternative splicing and environmental adaptation in wild house mice. Heredity 132:133–141.

Marden JH. 2008. Quantitative and evolutionary biology of alternative splicing: how changing the mix of alternative transcripts affects phenotypic plasticity and reaction norms. Heredity (Edinb) 100:111–120.

Martin Anduaga A, Evantal N, Patop IL, Bartok O, Weiss R, Kadener S. 2019. Thermosensitive alternative splicing senses and mediates temperature adaptation in Drosophila. Ramaswami M, Calabrese RL, editors. eLife 8:e44642.

McGirr JA, Martin CH. 2018. Parallel evolution of gene expression between trophic specialists despite divergent genotypes and morphologies. Evol Lett 2:62–75.

McManus CJ, Coolon JD, Eipper-Mains J, Wittkopp PJ, Graveley BR. 2014. Evolution of splicing regulatory networks in Drosophila. Genome Res. 24:786–796.

von Mering C, Jensen LJ, Snel B, Hooper SD, Krupp M, Foglierini M, Jouffre N, Huynen MA, Bork P. 2005. STRING: known and predicted protein–protein associations, integrated and transferred across organisms. Nucleic Acids Research 33:D433–D437.

Merkin J, Russell C, Chen P, Burge CB. 2012. Evolutionary dynamics of gene and isoform regulation in Mammalian tissues. Science 338:1593–1599.

Molière S, Jaulin A, Tomasetto C-L, Dali-Youcef N. 2023. Roles of Matrix Metalloproteinases and Their Natural Inhibitors in Metabolism: Insights into Health and Disease. International Journal of Molecular Sciences 24:10649.

Nomura K, Kinoshita S, Mizusaki N, Senga Y, Sasaki T, Kitamura T, Sakaue H, Emi A, Hosooka T, Matsuo M, et al. 2024. Adaptive gene expression of alternative splicing variants of PGC-1α regulates whole-body energy metabolism. Molecular Metabolism 86:101968.

O’Brown NM, Summers BR, Jones FC, Brady SD, Kingsley DM. 2015. A recurrent regulatory change underlying altered expression and Wnt response of the stickleback armor plates gene EDA. eLife 4:e05290.

Ostacolo K, López García de Lomana A, Larat C, Hjaltalin V, Holm KY, Hlynsdóttir SS, Soucheray M, Sooman L, Rolfsson O, Krogan NJ, et al. 2024. ATG7(2) Interacts With Metabolic Proteins and Regulates Central Energy Metabolism. Traffic 25:e12933.

Pan Q, Shai O, Lee LJ, Frey BJ, Blencowe BJ. 2008. Deep surveying of alternative splicing complexity in the human transcriptome by high-throughput sequencing. Nat Genet 40:1413–1415.

Papatheodorou I, Fonseca NA, Keays M, Tang YA, Barrera E, Bazant W, Burke M, Füllgrabe A, Fuentes AM-P, George N, et al. 2018. Expression Atlas: gene and protein expression across multiple studies and organisms. Nucleic Acids Res 46:D246–D251.

Pendás AM, Folgueras,Alicia R., Llano,Elena, Caterina,John, Frerard, Françoise, Rodríguez, Francisco, Astudillo, Aurora, Noël, Agnès, Birkedal-Hansen, Henning, and López-Otín C. 2004. Diet-Induced Obesity and Reduced Skin Cancer Susceptibility in Matrix Metalloproteinase 19-Deficient Mice. Molecular and Cellular Biology 24:5304–5313.

Phifer-Rixey M, Bi K, Ferris KG, Sheehan MJ, Lin D, Mack KL, Keeble SM, Suzuki TA, Good JM, Nachman MW. 2018. The genomic basis of environmental adaptation in house mice. PLOS Genetics 14:e1007672.

Price TD, Qvarnström A, Irwin DE. 2003. The role of phenotypic plasticity in driving genetic evolution. Proceedings of the Royal Society of London. Series B: Biological Sciences 270:1433–1440.

Qi T, Wu Y, Fang H, Zhang F, Liu S, Zeng J, Yang J. 2022. Genetic control of RNA splicing and its distinct role in complex trait variation. Nat Genet 54:1355–1363.

Ren G, Bhatnagar S, Young ME, Lee T, Kim J. 2025. Endothelial autophagy-related gene 7 contributes to high fat diet-induced obesity. Molecular Metabolism 93:102099.

Rogers TF, Palmer DH, Wright AE. 2020. Sex-Specific Selection Drives the Evolution of Alternative Splicing in Birds. Mol Biol Evol 38:519–530.

Schaefke B, Sun W, Li Y-S, Fang L, Chen W. 2018. The evolution of posttranscriptional regulation. WIREs RNA 9:e1485.

Scheiner SM. 1993. Genetics and Evolution of Phenotypic Plasticity. Annual Review of Ecology, Evolution, and Systematics 24:35–68.

Shen S, Park JW, Lu Z, Lin L, Henry MD, Wu YN, Zhou Q, Xing Y. 2014. rMATS: Robust and flexible detection of differential alternative splicing from replicate RNA-Seq data. Proceedings of the National Academy of Sciences 111:E5593–E5601.

Shpak M, Lawrence KN, Pool JE. 2025. The Precision and Power of Population Branch Statistics in Identifying the Genomic Signatures of Local Adaptation. Genome Biology and Evolution 17:evaf080.

Siddiq MA, Duveau F, Wittkopp PJ. 2024. Plasticity and environment-specific relationships between gene expression and fitness in Saccharomyces cerevisiae. Nat Ecol Evol 8:2184–2194.

Signor SA, Nuzhdin SV. 2018. The evolution of gene expression in cis and trans. Trends Genet 34:532–544.

Singh P, Ahi EP. 2022. The importance of alternative splicing in adaptive evolution. Molecular Ecology 31:1928–1938.

Singh P, Börger C, More H, Sturmbauer C. 2017. The Role of Alternative Splicing and Differential Gene Expression in Cichlid Adaptive Radiation. Genome Biology and Evolution 9:2764–2781.

Smith CCR, Tittes S, Mendieta JP, Collier-zans E, Rowe HC, Rieseberg LH, Kane NC. 2018. Genetics of alternative splicing evolution during sunflower domestication. Proceedings of the National Academy of Sciences 115:6768–6773.

Sommer RJ. 2020. Phenotypic Plasticity: From Theory and Genetics to Current and Future Challenges. Genetics 215:1–13.

Stern DL, Orgogozo V. 2008. The Loci of Evolution: How Predictable Is Genetic Evolution? Evolution 62:2155–2177.

Steward RA, de Jong MA, Oostra V, Wheat CW. 2022. Alternative splicing in seasonal plasticity and the potential for adaptation to environmental change. Nat Commun 13:755.

Szklarczyk D, Kirsch R, Koutrouli M, Nastou K, Mehryary F, Hachilif R, Gable AL, Fang T, Doncheva NT, Pyysalo S, et al. 2023. The STRING database in 2023: protein-protein association networks and functional enrichment analyses for any sequenced genome of interest. Nucleic Acids Res 51:D638–D646.

Thomas PD, Kejariwal A, Campbell MJ, Mi H, Diemer K, Guo N, Ladunga I, Ulitsky-Lazareva B, Muruganujan A, Rabkin S, et al. 2003. PANTHER: a browsable database of gene products organized by biological function, using curated protein family and subfamily classification. Nucleic Acids Res 31:334–341.

Tian L, Jabbari JS, Thijssen R, Gouil Q, Amarasinghe SL, Voogd O, Kariyawasam H, Du MRM, Schuster J, Wang C, et al. 2021. Comprehensive characterization of single-cell full-length isoforms in human and mouse with long-read sequencing. Genome Biology 22:310.

Tovar-Corona JM, Castillo-Morales A, Chen L, Olds BP, Clark JM, Reynolds SE, Pittendrigh BR, Feil EJ, Urrutia AO. 2015. Alternative Splice in Alternative Lice. Mol Biol Evol 32:2749–2759.

Vernia S, Edwards YJ, Han MS, Cavanagh-Kyros J, Barrett T, Kim JK, Davis RJ. 2016. An alternative splicing program promotes adipose tissue thermogenesis.Blencowe BJ, editor. eLife 5:e17672.

Verta J-P, Jacobs A. 2022. The role of alternative splicing in adaptation and evolution. Trends in Ecology & Evolution 37:299–308.

Verta J-P, Jones FC. 2019. Predominance of cis-regulatory changes in parallel expression divergence of sticklebacks.de Meaux J, Tautz D, editors. eLife 8:e43785.

Walter GM, Clark J, Cristaudo A, Terranova D, Nevado B, Catara S, Paunov M, Velikova V, Filatov D, Cozzolino S, et al. 2022. Adaptive divergence generates distinct plastic responses in two closely related Senecio species. Evolution 76:1229–1245.

Wang X, Yang M, Ren D, Terzaghi W, Deng X-W, He G. 2019. Cis-regulated alternative splicing divergence and its potential contribution to environmental responses in Arabidopsis. The Plant Journal 97:555–570.

Wang Y, Xie Z, Kutschera E, Adams JI, Kadash-Edmondson KE, Xing Y. 2024. rMATS-turbo: an efficient and flexible computational tool for alternative splicing analysis of large-scale RNA-seq data. Nat Protoc 19:1083–1104.

Watanabe K, Stringer S, Frei O, Umićević Mirkov M, de Leeuw C, Polderman TJC, van der Sluis S, Andreassen OA, Neale BM, Posthuma D. 2019. A global overview of pleiotropy and genetic architecture in complex traits. Nat Genet 51:1339–1348.

Wei W, Chen Q, Liu M, Sheng Y, OuYang Q, Feng W, Yang X, Ding L, Su S, Zhang J, et al. 2022. TRIM24 is an insulin-responsive regulator of P-bodies. Nat Commun 13:3972.

Weng M-P, Liao B-Y. 2017. modPhEA: model organism Phenotype Enrichment Analysis of eukaryotic gene sets. Bioinformatics 33:3505–3507.

West-Eberhard, M.J. 2003. Developmental Plasticity and Evolution. Oxford University Press Available from: https://academic.oup.com/book/40908

Wittkopp PJ, Haerum BK, Clark AG. 2004. Evolutionary changes in cis and trans gene regulation. Nature 430:85–88.

Wittkopp PJ, Kalay G. 2011. Cis-regulatory elements: molecular mechanisms and evolutionary processes underlying divergence. Nat Rev Genet 13:59–69.

Wright CJ, Smith CWJ, Jiggins CD. 2022. Alternative splicing as a source of phenotypic diversity. Nat Rev Genet 23:697–710.

Wright KM, Deighan AG, Di Francesco A, Freund A, Jojic V, Churchill GA, Raj A. 2022. Age and diet shape the genetic architecture of body weight in diversity outbred mice. Hägg S, Barkai N, Dahl A, Boud A, editors. eLife 11:e64329.

Yi X, Liang Y, Huerta-Sanchez E, Jin X, Cuo ZXP, Pool JE, Xu X, Jiang H, Vinckenbosch N, Korneliussen TS, et al. 2010. Sequencing of 50 Human Exomes Reveals Adaptation to High Altitude. Science 329:75–78.

Zhang Y, Goldman S, Baerga R, Zhao Y, Komatsu M, Jin S. 2009. Adipose-specific deletion of autophagy-related gene 7 (atg7) in mice reveals a role in adipogenesis. Proceedings of the National Academy of Sciences 106:19860–19865.

