## Supplemental Figures for "Alternative splicing contributes to plasticity and regulatory divergence in locally adapted house mice from the Americas"

### Supplementary Figures

**Figure S1.** Overlap between sex-biased genes across strains on a standard diet (top) and a high-fat diet (bottom).

STANDARD DIET

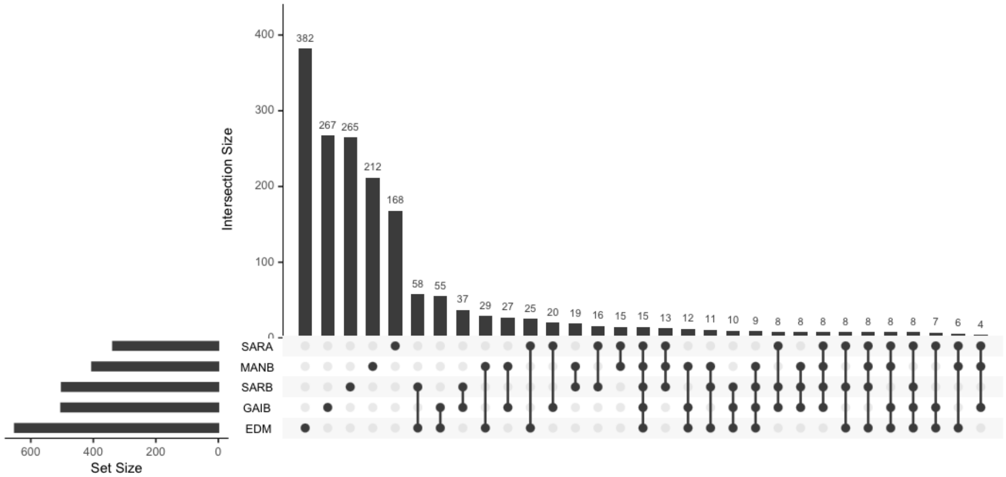


HF DIET

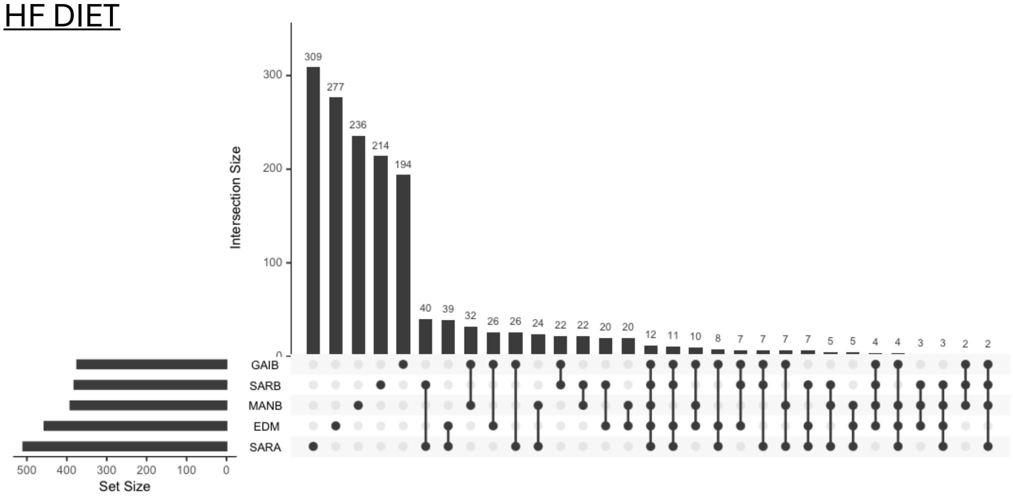

**Figure S2.** Features of differentially expressed genes [DEG] and differentially spliced genes [DSG] related to diet. **(A)** DEGs were associated with more protein-protein interactions [PPI] than DSGs. (**B**) There was not a significant difference in the number of GO Biological process terms between DSGs and DEGs. (**C**) DSGs were expressed across more tissue-developmental timepoints compared to DEGs.

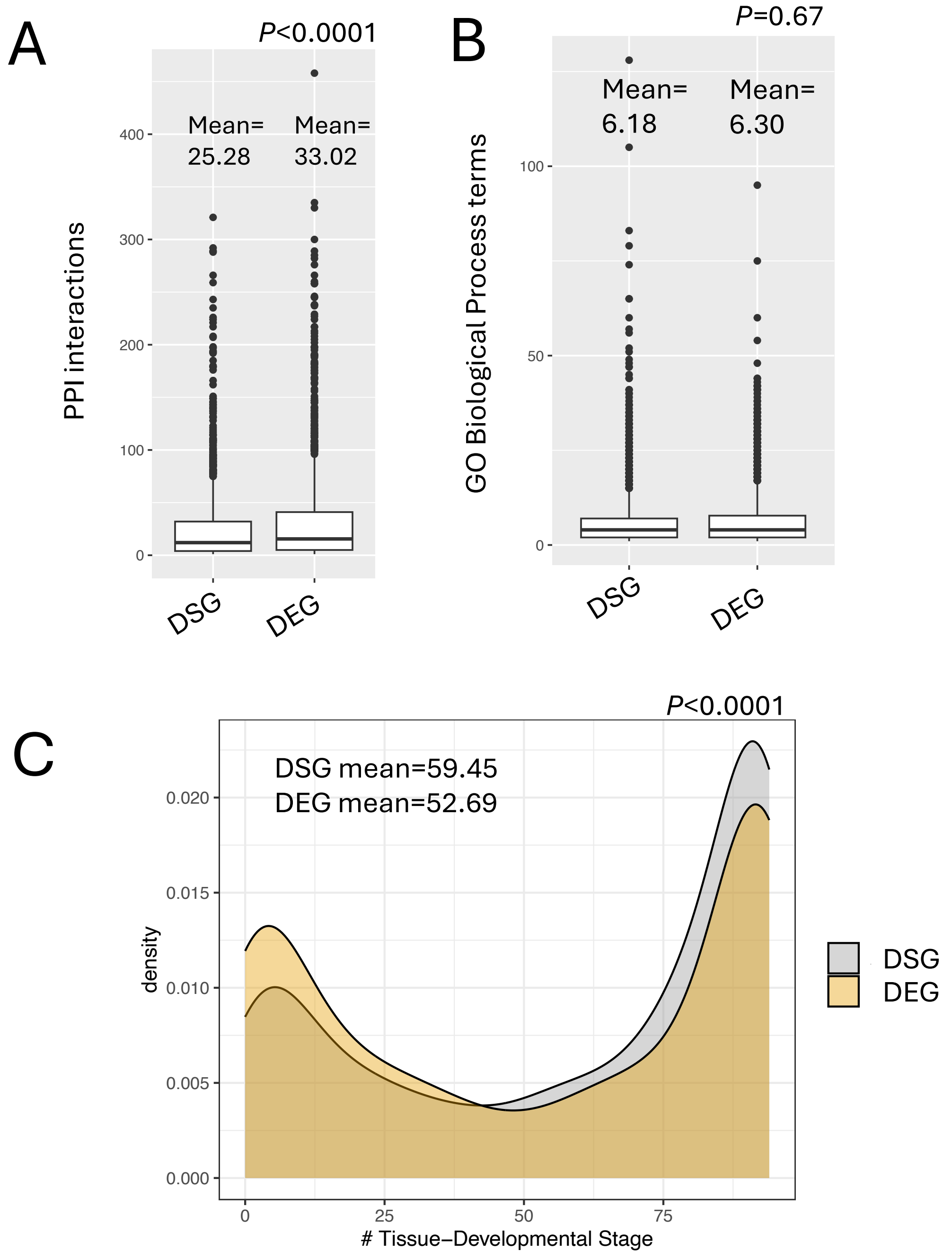
