## Supplemental Tables for "Alternative splicing contributes to plasticity and regulatory divergence in locally adapted house mice from the Americas"

### Supplementary Tables

**Table S1**. Number of differential splicing events between strains by category.

| Splice event | Unfiltered | Filtered by ΔPSI threshold |
|---|---|---|
| Alternative 5’ splice site | 799 | 412 |
| Alternative 3’ splice site | 1,133 | 522 |
| Mutually exclusive exons | 3,550 | 1,548 |
| Skipped exon | 8,247 | 4,165 |
| Retained intron | 1,063 | 340 |

**Table S2**. Differential alternative splicing events associated with diet identified for each strain and sex.

| Genotype | Skipped Exon | Alt. 5' splice site | Alt. 3 splice site | Mutually exclusive exons | Retained Intron |
|---|---|---|---|---|---|
| MANB, M | 191 | 49 | 35 | 37 | 27 |
| MANB, F | 235 | 23 | 64 | 25 | 27 |
| SARA, M | 209 | 20 | 45 | 13 | 46 |
| SARA, F | 237 | 20 | 47 | 120 | 26 |
| SARB, M | 217 | 25 | 50 | 18 | 24 |
| SARB, F | 242 | 21 | 24 | 46 | 62 |
| EDME, F | 287 | 48 | 60 | 43 | 230 |
| EDME, M | 199 | 26 | 24 | 24 | 20 |
| GAIB, F | 263 | 21 | 36 | 38 | 95 |
| GAIB, M | 221 | 29 | 24 | 37 | 21 |

**Table S3**. Genes with sex-biased alternative splicing identified for each strain on a high-fat versus standard diet.

| Strain | Standard | High-fat | Shared |
|---|---|---|---|
| BZ | 404 | 391 | 84 |
| EDME | 651 | 456 | 131 |
| FL | 503 | 374 | 90 |
| SARA | 337 | 509 | 89 |
| SARB | 501 | 381 | 97 |

**Table S4**. Genes with sex-biased expression across all strains under on either a high-fat or standard diet.

| Gene Symbol | Ensembl ID | Condition |
|---|---|---|
| *Zbtb20* | ENSMUSG00000022708 | Standard |
| *Xiap* | ENSMUSG00000025860 | Standard |
| *Scp2* | ENSMUSG00000028603 | Standard |
| *Sult2a8* | ENSMUSG00000030378 | Standard |
| *Ccnd3* | ENSMUSG00000034165 | Standard |
| *AI182371* | ENSMUSG00000035875 | Standard |
| *Mat1a* | ENSMUSG00000037798 | Standard |
| *Rpl22l1* | ENSMUSG00000039221 | Standard |
| *Kyat1* | ENSMUSG00000039648 | Standard |
| *Gas5* | ENSMUSG00000053332 | Standard |
| *Cyp3a59* | ENSMUSG00000061292 | Standard |
| *Selenbp2* | ENSMUSG00000068877 | Standard |
| *Gm11789* | ENSMUSG00000084983 | Standard |
| *Gm20319* | ENSMUSG00000092545 | Standard |
| *Jpx* | ENSMUSG00000097571 | Standard |
| *Ndrg2* | ENSMUSG00000004558 | High-fat |
| *C4a* | ENSMUSG00000015451 | High-fat |
| *Zbtb20* | ENSMUSG00000022708 | High-fat |
| *Serpinc1* | ENSMUSG00000026715 | High-fat |
| *Scp2* | ENSMUSG00000028603 | High-fat |
| *Alb* | ENSMUSG00000029368 | High-fat |
| *Slco1a4* | ENSMUSG00000030237 | High-fat |
| *Sult2a8* | ENSMUSG00000030378 | High-fat |
| *AI182371* | ENSMUSG00000035875 | High-fat |
| *Rpl22l1* | ENSMUSG00000039221 | High-fat |
| *Gm20544* | ENSMUSG00000092569 | High-fat |
| *BC024386* | ENSMUSG00000109628 | High-fat |

**Table S5**. Overlap between differentially spliced and differentially expressed genes binned by expression level. Mean normalized expression is given in parentheses for each bin.

| | | **Differential expression** | **No Differential expression** |
|---|---|---|---|
| **Bin 1** (44) | **Differential splicing** | 46 | 83 |
| | **No differential splicing** | 848 | 1706 |
| **Bin 2** (183) | **Differential splicing** | 105 | 168 |
| | **No differential splicing** | 761 | 1650 |
| **Bin 3** (536) | **Differential splicing** | 129 | 283 |
| | **No differential splicing** | 563 | 1708 |
| **Bin 4** (1,273) | **Differential splicing** | 131 | 338 |
| | **No differential splicing** | 479 | 1735 |
| **Bin 5** (4,483) | **Differential splicing** | 199 | 419 |
| | **No differential splicing** | 469 | 1596 |

**Table S6.** Genes with evidence for allele-specific alternative splicing on each diet.

| | | | **Splice sites** | **Splice sites, ΔPSI>0.1** | **Genes** | **Genes, ΔPSI>0.1** |
|---|---|---|---|---|---|---|
| **SARBxMANB** | M | High-fat | 384 | 242 | 272 | 175 |
| | | Standard | 454 | 255 | 322 | 187 |
| | F | High-fat | 484 | 283 | 341 | 201 |
| | | Standard | 405 | 241 | 289 | 184 |
| **GAIBxSARA** | M | High-fat | 375 | 202 | 274 | 160 |
| | | Standard | 384 | 232 | 282 | 178 |
| | F | High-fat | 403 | 214 | 291 | 174 |
| | | Standard | 405 | 241 | 289 | 184 |

**Table S7.** *Cis*- vs. *trans*- changes contributing to divergence in alternative splicing (A3SS: alternative 3' splice site; A5SS: alternative 5' splice site; RI: retained intron; SE: skipped exon)

| Cross | Event type | Total examined | *Cis* only | *Trans* only |
|---|---|---|---|---|
| **SARBxMANB, male** | RI | 177 | 14 | 0 |
| | SE | 3885 | 159 | 29 |
| | A5SS | 108 | 5 | 0 |
| | A3SS | 180 | 4 | 0 |
| **SARBxMANB, female** | RI | 156 | 6 | 0 |
| | SE | 4177 | 167 | 27 |
| | A5SS | 131 | 3 | 1 |
| | A3SS | 186 | 2 | 2 |
| **GAIBxSARA, male** | RI | 122 | 4 | 4 |
| | SE | 3450 | 128 | 18 |
| | A5SS | 105 | 0 | 0 |
| | A3SS | 131 | 4 | 4 |
| **GAIBxSARA, female** | RI | 185 | 5 | 0 |
| | SE | 4100 | 140 | 17 |
| | A5SS | 112 | 4 | 0 |
| | A3SS | 202 | 8 | 2 |

**Table S8**. The number of differentially spliced genes that also co-localize with outlier loci from selection analyses for each focal population.

| Focal Population | Co-localizing genes | Co-localizing genes, 10% ΔPSI cut-off |
|---:|---:|---:|
| Florida, USA | 54 | 28 |
| Alberta, Canada | 47 | 28 |
| Amazonas, Brazil | 74 | 43 |
| Northeastern USA | 74 | 46 |

**Table S9.** Enrichments for outliers with evidence for differential splicing.

| Focal population | Phenotype Term related to metabolism/homeostasis | *q*-value |
|---|---|---|
| Northeastern USA | homeostasis/metabolism phenotype | 0.002 |
| Northeastern USA | abnormal homeostasis | 0.004 |
| Manaus, Brazil | abnormal postnatal growth/weight/body size | 0.001 |
| Manaus, Brazil | homeostasis/metabolism phenotype | 0.002 |
| Manaus, Brazil | abnormal body size | 0.002 |
| Manaus, Brazil | abnormal homeostasis | 0.007 |
| Manaus, Brazil | decreased body size | 0.012 |
| Manaus, Brazil | abnormal body weight | 0.012 |
| Manaus, Brazil | abnormal body composition | 0.016 |
| Manaus, Brazil | adipose tissue phenotype | 0.036 |
| Manaus, Brazil | abnormal adipose tissue morphology | 0.047 |
| Manaus, Brazil | homeostasis/metabolism phenotype | 0.006 |
| Manaus, Brazil | growth/size/body region phenotype | 0.013 |
| Manaus, Brazil | abnormal homeostasis | 0.021 |
| Edmonton, Canada | growth/size/body region phenotype | 0.009 |
| Edmonton, Canada | abnormal postnatal growth/weight/body size | 0.01 |
| Edmonton, Canada | abnormal body size | 0.022 |
| Edmonton, Canada | homeostasis/metabolism phenotype | 0.049 |
| Florida, USA | homeostasis/metabolism phenotype | 0.006 |
| Florida, USA | growth/size/body region phenotype | 0.013 |
| Florida, USA | abnormal homeostasis | 0.021 |

**Table S10.** Splicing-related genomic outliers.

| Ensembl ID | Gene symbol | Focal Population |
|---|---|---|
| ENSMUSG00000001089 | *Luzp1* | Florida |
| ENSMUSG00000001120 | *Pcbp3* | Amazonas |
| ENSMUSG00000001247 | *Lsr* | Florida |
| ENSMUSG00000001376 | *Vps50* | Northeastern USA |
| ENSMUSG00000001998 | *Ap4e1* | Northeastern USA |
| ENSMUSG00000002846 | *Timmdc1* | Florida |
| ENSMUSG00000003273 | *Car11* | Florida |
| ENSMUSG00000003363 | *Pld3* | Amazonas |
| ENSMUSG00000003992 | *Ssbp2* | Alberta |
| ENSMUSG00000004056 | *Akt2* | Amazonas |
| ENSMUSG00000004266 | *Ptpn6* | Northeastern USA |
| ENSMUSG00000004565 | *Pnpla6* | Florida |
| ENSMUSG00000004789 | *Dlst* | Northeastern USA |
| ENSMUSG00000005102 | *Eif2ak4* | Amazonas |
| ENSMUSG00000006673 | *Qrich1* | Amazonas |
| ENSMUSG00000014030 | *Pax5* | Northeastern USA |
| ENSMUSG00000014074 | *Rnf168* | Northeastern USA |
| ENSMUSG00000014850 | *Msh3* | Alberta |
| ENSMUSG00000015143 | *Actn1* | Northeastern USA |
| ENSMUSG00000015968 | *Cacna1d* | Florida |
| ENSMUSG00000016918 | *Sulf1* | Northeastern USA |
| ENSMUSG00000017723 | *Wfdc2* | Amazonas |
| ENSMUSG00000017978 | *Cadps2* | Amazonas |
| ENSMUSG00000018167 | *Stard3* | Northeastern USA |
| ENSMUSG00000018334 | *Ksr1* | Northeastern USA |
| ENSMUSG00000018411 | *Mapt* | Alberta |
| ENSMUSG00000018800 | *Abca5* | Florida |
| ENSMUSG00000018906 | *P4ha2* | Northeastern USA |
| ENSMUSG00000019989 | *Enpp3* | Alberta |
| ENSMUSG00000020153 | *Ndufs7* | Northeastern USA |
| ENSMUSG00000020220 | *Vps13d* | Northeastern USA |
| ENSMUSG00000020284 | *Cfap410* | Northeastern USA |
| ENSMUSG00000020654 | *Adcy3* | Amazonas |
| ENSMUSG00000020709 | *Adap2* | Florida |
| ENSMUSG00000020954 | *Strn3* | Northeastern USA |
| ENSMUSG00000021112 | *Pals1* | Florida |
| ENSMUSG00000021140 | *Pcnx* | Alberta |

| | | |
|---|---|---|
| ENSMUSG00000021140 | *Pcnx* | Northeastern USA |
| ENSMUSG00000021180 | *Rps6ka5* | Northeastern USA |
| ENSMUSG00000021266 | *Wars1* | Northeastern USA |
| ENSMUSG00000021375 | *Kif13a* | Florida |
| ENSMUSG00000021519 | *Mterf3* | Florida |
| ENSMUSG00000021576 | *Pdcd6* | Florida |
| ENSMUSG00000021619 | *Atg10* | Florida |
| ENSMUSG00000021759 | *Plpp1* | Florida |
| ENSMUSG00000021759 | *Plpp1* | Northeastern USA |
| ENSMUSG00000021990 | *Spata13* | Florida |
| ENSMUSG00000022237 | *Ankrd33b* | Northeastern USA |
| ENSMUSG00000022629 | *Kif21a* | Amazonas |
| ENSMUSG00000022748 | *Cmss1* | Amazonas |
| ENSMUSG00000022853 | *Ehhadh* | Northeastern USA |
| ENSMUSG00000023345 | *Poc1a* | Amazonas |
| ENSMUSG00000023353 | *Agap3* | Northeastern USA |
| ENSMUSG00000023963 | *Cyp39a1* | Amazonas |
| ENSMUSG00000024044 | *Epb41l3* | Amazonas |
| ENSMUSG00000024104 | *Washc2* | Northeastern USA |
| ENSMUSG00000024160 | *Spsb3* | Alberta |
| ENSMUSG00000024163 | *Mapk8ip3* | Alberta |
| ENSMUSG00000024238 | *Zeb1* | Alberta |
| ENSMUSG00000024413 | *Npc1* | Florida |
| ENSMUSG00000024589 | *Nedd4l* | Northeastern USA |
| ENSMUSG00000024740 | *Ddb1* | Alberta |
| ENSMUSG00000024773 | *Atg2a* | Alberta |
| ENSMUSG00000024948 | *Map4k2* | Alberta |
| ENSMUSG00000025006 | *Sorbs1* | Alberta |
| ENSMUSG00000025064 | *Col17a1* | Alberta |
| ENSMUSG00000025085 | *Ablim1* | Amazonas |
| ENSMUSG00000025185 | *Loxl4* | Alberta |
| ENSMUSG00000025199 | *Chuk* | Amazonas |
| ENSMUSG00000025234 | *Arih1* | Florida |
| ENSMUSG00000025355 | *Mmp19* | Amazonas |
| ENSMUSG00000025404 | *R3hdm2* | Northeastern USA |
| ENSMUSG00000025512 | *Chid1* | Northeastern USA |
| ENSMUSG00000025871 | *4833439L19Rik* | Northeastern USA |
| ENSMUSG00000025911 | *Adhfe1* | Amazonas |
| ENSMUSG00000025911 | *Adhfe1* | Florida |
| ENSMUSG00000025958 | *Creb1* | Alberta |

| | | |
|---|---|---|
| ENSMUSG00000026048 | *Ercc5* | Amazonas |
| ENSMUSG00000026049 | *Tex30* | Amazonas |
| ENSMUSG00000026113 | *Inpp4a* | Florida |
| ENSMUSG00000026463 | *Atp2b4* | Florida |
| ENSMUSG00000026614 | *Slc30a10* | Amazonas |
| ENSMUSG00000026655 | *Fam107b* | Alberta |
| ENSMUSG00000026657 | *Frmd4a* | Florida |
| ENSMUSG00000026674 | *Ddr2* | Northeastern USA |
| ENSMUSG00000026723 | *Trdmt1* | Florida |
| ENSMUSG00000026727 | *Rsu1* | Alberta |
| ENSMUSG00000026778 | *Prkcq* | Northeastern USA |
| ENSMUSG00000026918 | *Brd3* | Northeastern USA |
| ENSMUSG00000026921 | *Egfl7* | Alberta |
| ENSMUSG00000026921 | *Egfl7* | Florida |
| ENSMUSG00000026921 | *Egfl7* | Northeastern USA |
| ENSMUSG00000026950 | *Neb* | Alberta |
| ENSMUSG00000027048 | *Abcb11* | Alberta |
| ENSMUSG00000027519 | *Rab22a* | Northeastern USA |
| ENSMUSG00000027531 | *Impa1* | Northeastern USA |
| ENSMUSG00000027556 | *Car1* | Northeastern USA |
| ENSMUSG00000027901 | *Dennd2d* | Florida |
| ENSMUSG00000028152 | *Tspan5* | Florida |
| ENSMUSG00000028329 | *Xpa* | Florida |
| ENSMUSG00000028399 | *Ptprd* | Amazonas |
| ENSMUSG00000028399 | *Ptprd* | Florida |
| ENSMUSG00000028433 | *Ubap2* | Northeastern USA |
| ENSMUSG00000028538 | *St3gal3* | Florida |
| ENSMUSG00000029095 | *Ablim2* | Northeastern USA |
| ENSMUSG00000029125 | *Stx18* | Amazonas |
| ENSMUSG00000029127 | *Zbtb49* | Alberta |
| ENSMUSG00000029446 | *Psph* | Amazonas |
| ENSMUSG00000029461 | *Fam168a* | Northeastern USA |
| ENSMUSG00000029695 | *Aass* | Amazonas |
| ENSMUSG00000029722 | *Agfg2* | Amazonas |
| ENSMUSG00000029826 | *Zc3hav1* | Northeastern USA |
| ENSMUSG00000029924 | *Slc37a3* | Northeastern USA |
| ENSMUSG00000030131 | *Mug2* | Northeastern USA |
| ENSMUSG00000030177 | *Ccdc77* | Northeastern USA |
| ENSMUSG00000030228 | *Pik3c2g* | Amazonas |
| ENSMUSG00000030302 | *Atp2b2* | Amazonas |

| | | |
|---|---|---|
| ENSMUSG00000030513 | *Pcsk6* | Northeastern USA |
| ENSMUSG00000030583 | *Sipa1l3* | Northeastern USA |
| ENSMUSG00000030760 | *Acer3* | Alberta |
| ENSMUSG00000030763 | *Lcmt1* | Amazonas |
| ENSMUSG00000030766 | *Arhgap17* | Amazonas |
| ENSMUSG00000030780 | *Rusf1* | Northeastern USA |
| ENSMUSG00000030834 | *Abcc6* | Amazonas |
| ENSMUSG00000030849 | *Fgfr2* | Alberta |
| ENSMUSG00000030849 | *Fgfr2* | Amazonas |
| ENSMUSG00000030849 | *Fgfr2* | Florida |
| ENSMUSG00000030852 | *Tacc2* | Alberta |
| ENSMUSG00000031511 | *Arhgef7* | Northeastern USA |
| ENSMUSG00000031533 | *Mrps31* | Florida |
| ENSMUSG00000031565 | *Fgfr1* | Amazonas |
| ENSMUSG00000031586 | *Rbpms* | Amazonas |
| ENSMUSG00000031816 | *Mthfsd* | Amazonas |
| ENSMUSG00000031822 | *Gse1* | Amazonas |
| ENSMUSG00000032058 | *Ppp2r1b* | Alberta |
| ENSMUSG00000032263 | *Bckdhb* | Northeastern USA |
| ENSMUSG00000032314 | *Etfa* | Northeastern USA |
| ENSMUSG00000032913 | *Lrig2* | Alberta |
| ENSMUSG00000033082 | *Clec1a* | Alberta |
| ENSMUSG00000033632 | *AW554918* | Florida |
| ENSMUSG00000034118 | *Tpst1* | Northeastern USA |
| ENSMUSG00000034211 | *Mrps17* | Amazonas |
| ENSMUSG00000034647 | *Ankrd12* | Alberta |
| ENSMUSG00000034647 | *Ankrd12* | Northeastern USA |
| ENSMUSG00000034774 | *Dsg1c* | Northeastern USA |
| ENSMUSG00000034826 | *Nup54* | Amazonas |
| ENSMUSG00000035069 | *Oma1* | Northeastern USA |
| ENSMUSG00000035270 | *Impg2* | Alberta |
| ENSMUSG00000035270 | *Impg2* | Florida |
| ENSMUSG00000035637 | *Grhpr* | Amazonas |
| ENSMUSG00000035778 | *Ggta1* | Amazonas |
| ENSMUSG00000036098 | *Myrf* | Amazonas |
| ENSMUSG00000036555 | *Iqce* | Florida |
| ENSMUSG00000036743 | *Psma8* | Amazonas |
| ENSMUSG00000036825 | *Ssx2ip* | Amazonas |
| ENSMUSG00000037210 | *Fam193a* | Amazonas |
| ENSMUSG00000037426 | *Depdc5* | Northeastern USA |

| | | |
|---|---|---|
| ENSMUSG00000037541 | *Shank2* | Amazonas |
| ENSMUSG00000038260 | *Trpm4* | Northeastern USA |
| ENSMUSG00000038591 | *Colec10* | Amazonas |
| ENSMUSG00000038658 | *Ric1* | Florida |
| ENSMUSG00000039099 | *Wdr93* | Alberta |
| ENSMUSG00000039099 | *Wdr93* | Northeastern USA |
| ENSMUSG00000039230 | *Tbcd* | Florida |
| ENSMUSG00000039456 | *Morc3* | Amazonas |
| ENSMUSG00000039473 | *Ubn1* | Northeastern USA |
| ENSMUSG00000039648 | *Kyat1* | Amazonas |
| ENSMUSG00000039662 | *Icmt* | Florida |
| ENSMUSG00000039842 | *Mcph1* | Alberta |
| ENSMUSG00000039879 | *Heca* | Northeastern USA |
| ENSMUSG00000039914 | *Coq10a* | Amazonas |
| ENSMUSG00000039934 | *Gsap* | Alberta |
| ENSMUSG00000040410 | *Fbxl4* | Amazonas |
| ENSMUSG00000040464 | *Gtpbp10* | Northeastern USA |
| ENSMUSG00000040481 | *Bptf* | Amazonas |
| ENSMUSG00000040548 | *Tex2* | Florida |
| ENSMUSG00000040550 | *Otud6b* | Amazonas |
| ENSMUSG00000040583 | *Cyp2b13* | Florida |
| ENSMUSG00000041225 | *Arhgap12* | Alberta |
| ENSMUSG00000041341 | *Atg2b* | Alberta |
| ENSMUSG00000041650 | *Pcca* | Florida |
| ENSMUSG00000041828 | *Abca8a* | Florida |
| ENSMUSG00000042520 | *Ubap2l* | Amazonas |
| ENSMUSG00000042790 | *Rnf214* | Alberta |
| ENSMUSG00000043284 | *Tmem11* | Amazonas |
| ENSMUSG00000043671 | *Dpy19l3* | Northeastern USA |
| ENSMUSG00000043987 | *Cep164* | Northeastern USA |
| ENSMUSG00000045078 | *Rnf216* | Northeastern USA |
| ENSMUSG00000045205 | *Dpy19l4* | Amazonas |
| ENSMUSG00000045576 | *St7l* | Amazonas |
| ENSMUSG00000045973 | *Slc25a51* | Florida |
| ENSMUSG00000046798 | *Cldn12* | Amazonas |
| ENSMUSG00000046798 | *Cldn12* | Northeastern USA |
| ENSMUSG00000046897 | *Zfp740* | Alberta |
| ENSMUSG00000049421 | *Zfp260* | Northeastern USA |
| ENSMUSG00000050541 | *Adra1b* | Alberta |
| ENSMUSG00000050549 | *Fam241a* | Amazonas |

| | | |
|---|---|---|
| ENSMUSG00000051359 | *Ncald* | Amazonas |
| ENSMUSG00000052407 | *Ccdc171* | Amazonas |
| ENSMUSG00000052889 | *Prkcb* | Alberta |
| ENSMUSG00000052889 | *Prkcb* | Amazonas |
| ENSMUSG00000052889 | *Prkcb* | Northeastern USA |
| ENSMUSG00000053279 | *Aldh1a1* | Northeastern USA |
| ENSMUSG00000054874 | *Pcnx3* | Amazonas |
| ENSMUSG00000054874 | *Pcnx3* | Florida |
| ENSMUSG00000055296 | *Tmem245* | Alberta |
| ENSMUSG00000055546 | *Timd4* | Northeastern USA |
| ENSMUSG00000055720 | *Ubl7* | Florida |
| ENSMUSG00000055884 | *Fancm* | Northeastern USA |
| ENSMUSG00000056035 | *Cyp3a11* | Alberta |
| ENSMUSG00000057530 | *Ece1* | Florida |
| ENSMUSG00000057596 | *Trim30d* | Florida |
| ENSMUSG00000057933 | *Gsta2* | Alberta |
| ENSMUSG00000058239 | *Usf2* | Florida |
| ENSMUSG00000059119 | *Nap1l4* | Northeastern USA |
| ENSMUSG00000059669 | *Taf1b* | Amazonas |
| ENSMUSG00000059824 | *Dbp* | Amazonas |
| ENSMUSG00000059824 | *Dbp* | Florida |
| ENSMUSG00000061950 | *Ppp4r1* | Northeastern USA |
| ENSMUSG00000062234 | *Gak* | Florida |
| ENSMUSG00000062646 | *Ganc* | Amazonas |
| ENSMUSG00000063253 | *Scoc* | Northeastern USA |
| ENSMUSG00000063382 | *Bcl9l* | Amazonas |
| ENSMUSG00000063450 | *Syne2* | Alberta |
| ENSMUSG00000063450 | *Syne2* | Northeastern USA |
| ENSMUSG00000064294 | *Aox3* | Florida |
| ENSMUSG00000066735 | *Vkorc1l1* | Northeastern USA |
| ENSMUSG00000066829 | *Zfp810* | Alberta |
| ENSMUSG00000068037 | *Mas1* | Northeastern USA |
| ENSMUSG00000070565 | *Rasal2* | Amazonas |
| ENSMUSG00000073436 | *Eme2* | Alberta |
| ENSMUSG00000073664 | *Nbeal1* | Amazonas |
| ENSMUSG00000074377 | *Sult2a4* | Northeastern USA |
| ENSMUSG00000075486 | *Commd6* | Amazonas |
| ENSMUSG00000078515 | *Ddi2* | Amazonas |
| ENSMUSG00000083282 | *Ctsf* | Florida |
| ENSMUSG00000090100 | *Ttbk2* | Amazonas |

| | | |
|---|---|---|
| ENSMUSG00000090124 | *Ugt1a7c* | Alberta |
| ENSMUSG00000090175 | *Ugt1a9* | Alberta |
| ENSMUSG00000091476 | *Catspere2* | Alberta |
| ENSMUSG00000091476 | *Catspere2* | Florida |
| ENSMUSG00000097195 | *Snhg5* | Florida |
| ENSMUSG00000097277 | *2900076A07Rik* | Amazonas |
| ENSMUSG00000097277 | *2900076A07Rik* | Florida |
| ENSMUSG00000097354 | *2310001H17Rik* | Northeastern USA |
| ENSMUSG00000098234 | *Snhg6* | Amazonas |
| ENSMUSG00000109644 | *0610005C13Rik* | Amazonas |
| ENSMUSG00000109644 | *0610005C13Rik* | Florida |
| ENSMUSG00000112006 | *Gm48633* | Amazonas |
